# Mimicking physiological stiffness or oxygen levels in vitro reorganizes mesenchymal stem cells machinery toward a more naïve phenotype

**DOI:** 10.1101/2024.06.11.598426

**Authors:** Inês Caramelo, Vera M. Mendes, Catarina Domingues, Sandra I. Anjo, Margarida Geraldo, Carla M. P. Cardoso, Mário Grãos, Bruno Manadas

**Affiliations:** CNC-Center for Neuroscience and Cell Biology, University of Coimbra, 3004-504 Coimbra, Portugal; PhD Programme in Experimental Biology and Biomedicine, Institute for Interdisciplinary Research (IIIUC), University of Coimbra, Casa Costa Alemão, 3030-789 Coimbra, Portugal; CIBB - Centre for Innovative Biomedicine and Biotechnology, University of Coimbra; Institute for Interdisciplinary Research, University of Coimbra (IIIUC), 3030-789 Coimbra, Portugal; Stemlab S.A. - Crioestaminal, 3060-197 Cantanhede, Portugal; Biocant, Technology Transfer Association, 3060-197 Cantanhede, Portugal

**Keywords:** Mesenchymal stem cells, Mechanomodulation, Physioxia, Proteomics

## Abstract

Mesenchymal stem cells (MSCs) offer a promising therapeutic potential for a wide variety of pathologies. However, obtaining minimal effective doses requires an extensive *in vitro* expansion, which compromises their stemness and therapeutic properties. The stiffness of the umbilical cord ranges between 2 and 5kPa, and the oxygen levels fluctuate from 2.4% to 3.8%, differing from the standard *in vitro* culture conditions where MSCs are exposed to the stiffness of the Petri dish (2-3 GPa) and near atmospheric oxygen levels (18.5% O_2_). Since MSCs can sense and respond to biomechanical and chemical characteristics of the microenvironment, it was hypothesized that expanding MSCs on 3kPa platforms – mechanomodulation – or at 5% O_2_ levels – physioxia – could potentially impact the cellular proteome of MSCs, for long (7-10 days) or short (48h) periods. Data analysis has unveiled that culturing MSCs on soft substrates for long periods promotes the expression of various proteins related to cell redox homeostasis, such as thioredoxins and peroxiredoxins. Conversely, culturing these cells during the same period but under low oxygen levels leads to an increase in chaperone machinery proteins, such as HSP90 or TRiC. These proteins can favor the clearance of misfolded proteins and telomerase maintenance processes, possibly preventing MSCs from being driven to a senescent phenotype. Although mechanomodulation and physioxia are two distinct stimuli, both converge in downregulating the expression of histones and several ribosomal subunits, possibly decreasing translational complexity, which could hypothetically favor a more naïve phenotype for MSCs. Interestingly, priming UC-MSCs (48h) leads to a differential expression of proteins of the extracellular matrix and histone subtypes. Understanding the role of these proteins in transducing environmental cues might provide insights into how conventional culture conditions significantly alter fundamental cellular processes and support the development of a more efficient protocol to expand and empower the therapeutic potential of MSCs. In the future, employing a combination of reduced stiffness and lower oxygen levels may present a promising strategic approach.

**Highlights:**

- Culturing MSCs on a soft substrate (3kPa) enhances the expression of antioxidant proteins, such as thioredoxins and peroxiredoxins
- Protein homeostasis is remodeled in MSCs cultured under physiological levels of oxygen (5% O_2_) through the differential expression of the chaperone machinery
- Lowering stiffness or oxygen levels during *in vitro* MSCs expansion decreases histones and ribosomal subunits expression, possibly favoring a more naïve phenotype

## Introduction

In recent years, mesenchymal stem cells (MSCs) have emerged as a promising therapeutic strategy for a wide variety of pathologies, from neurological to cartilage disorders (1-3). When infused, these cells are chemoattracted to the site of injury, where they can exert their beneficial effects. They can penetrate the damaged tissue and act through paracrine signaling via secretion of soluble factors and/or extracellular vesicles that modulate inflammation, apoptosis and/or promote tissue regeneration (4-8).

Despite the promising therapeutic effects, the minimal effective dose for intravenous delivery ranges from 70 to 190 million cells/patient/dose, demanding an extensive *in vitro* expansion of MSCs (9). In fact, the increasing number of passages has been described as hindering their differentiation potential and immunomodulatory properties, potentially compromising their therapeutic properties (10, 11). Additionally, genomic alterations may occur during expansion, although they have not yet been associated with malignant transformations (12). Consequently, there is a need to develop new strategies to minimize the impact of *in vitro* expansion and enhance the therapeutical potential of MSCs.

Cells in living tissues, including MSCs, can sense and respond to the biophysical and biomechanical properties of the surrounding environment, like stiffness, which affects various biological processes (13). Specifically, protein complexes at the cell membrane, such as focal adhesions, act as mechanosensors. They convert mechanical cues of the extracellular matrix into biochemical signals by activating intracellular signaling pathways, a process called mechanotransduction. Additionally, since focal adhesions are connected to cytoskeleton proteins, this process affects cell morphology and allows signals to be transmitted directly to the nucleoskeleton, thereby modulating gene expression, and determining the fate of stem cells (13-15). Moreover, cells possess the ability to sense oxygen levels within each organ, which is known to impact cell proliferation and differentiation (16). In high oxygen environments, elevated levels of reactive oxygen species (ROS) stabilize the transcription factor Nrf2, prompting its translocation to the nucleus and subsequent upregulation of cytoprotective gene expression (17, 18). Conversely, the stabilization of HIF-1/2α, an intracellular O_2_ sensor, occurs in low oxygen tension environments. In such conditions, HIF-1/2α translocates to the nucleus, dimerizes with HIF-β, and regulates various target genes involved in cell stemness and self-renewal, redox homeostasis, among others (18, 19). Collectively, these findings suggest that both mechanical cues and oxygen levels play a pivotal role in determining cell behavior.

MSCs can be obtained from different sources (8). Particularly, umbilical cord MSCs (UC-MSCs) isolated from the Wharton’s Jelly exhibit faster proliferation rates compared to other adult MSCs, and share similarities with fetal-derived MSCs (20). Wharton’s jelly stiffness, quantified by the Young’s modulus (E), has been reported to range from 2 to 5 kPa (21). Moreover, the oxygen tension in the umbilical cord blood varies from around 2.4% to 3.8% (22). Hence, it is important to note that *in vitro* cell culture conditions differ significantly from the physiological characteristics of the tissue of origin from where MSCs are isolated. Under standard culture conditions, MSCs are exposed to approximately 18.5% O_2_ in an incubator and encounter the stiffness of a Petri dish, which ranges from 2-3 GPa (22, 23). In this study, the cellular proteome of MSCs cultured for long (7-10 days) and short periods (48h) under conditions that mimic a physiological environment (3kPa or 5% O_2_) was compared to standard culture conditions. The identification of differentially expressed proteins involved in the redox and protein homeostasis, as well as extracellular matrix proteins, suggests that these approaches might impact MSCs stemness and, consequently, their therapeutic potential.

## Methods

### Cell culture

#### Isolation and expansion of UC-MSCs

All procedures were approved by the Faculty of Medicine Ethics Committee, University of Coimbra, Portugal (ref. CE-075/2019) and were previously reported by our team (24). Succinctly, human MSCs were isolated from the UC (Wharton’s Jelly) and expanded with proliferation medium (Minimum essential Medium-α (MEM-α) (Gibco™) supplemented with 10% (v/v) MSC-qualified fetal bovine serum (FBS) (Hyclone, GE Healthcare) or 5% (v/v) fibrinogen depleted human platelet lysate (HPL) (UltraGRO™, Helios) and antibiotics: 100 U/ml of Penicillin, 100 μg/ml Streptomycin and 2.5 μg/ml Amphotericin B or Antibiotic-Antimycotic (all from Gibco™)). When sub-confluent, cells were washed with Phosphate Buffered Saline (PBS), detached by the addition of 0.05% Trypsin-EDTA solution (Gibco™) for five minutes at 37°C on a humidified incubator, homogenized and centrifuged for 5 minutes at 290×*g* at room temperature. Supernatant was discarded, the pellet was resuspended and MSCs were re-plated at the desired cell density.

#### Expansion of UC-MSCs on soft substrate (mechanomodulation)

Soft polydimethylsiloxane (PDMS) cell culture substrates were treated and UC-MSCs were expanded as previously described by Domingues *et. al* (24) (Figure 1A). Data concerning long exposure to soft culture conditions was previously published (24) and reanalyzed for the purpose of the present work. For priming experiments, when cell culture reached 80-90% confluence, cells were split 1:3 until P3 at standard conditions (Figure 4A). At P4, UC-MSCs were seeded at 10,000 cells/cm^2^ (P4) on PDMS platforms or TCP dishes and left for 48 hours.

**Figure 1.**
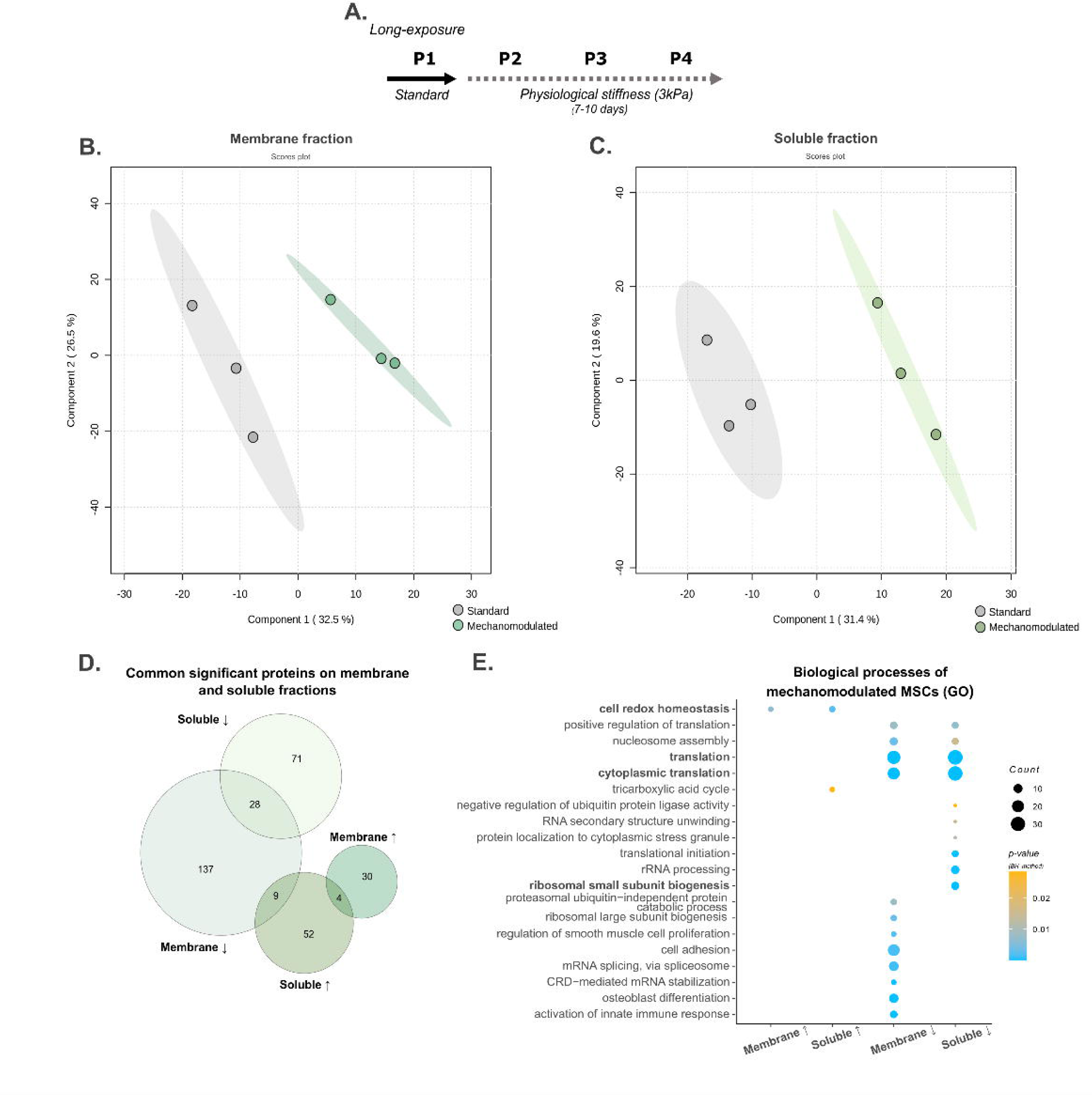
*Expanding MSCs on low stiffness platforms differentially modulate cell redox homeostasis and translation processes.* MSCs were expanded for 7 to 10 days (from P2 to P4) on 3kPa PDMS substrates, and the cellular proteome was characterized by DDA and DIA LC-MS/MS proteomics (A). Two PLS-DA models were generated individually using all proteins quantified with confidence for membrane (dark green) (B) and soluble (light green) (C) fractions; the standard condition is represented in grey. In total, 65 and 34 proteins were found to be up- (↑) regulated on the soluble and membrane fraction, respectively, and 99 and 165 to be down- (↓) regulated on the soluble and membrane fraction. Common proteins across fractions are represented on the Venn diagram (D). Proteins of interest underwent GO analysis, and the significantly enriched biological processes are depicted in (E). Abbreviations: Benjamini-Hochberg method (BH).

#### Expansion of UC-MSCs under controlled oxygen levels (physioxia)

For long exposure experiments, UC-MSCs were initially seeded at 4000 cells/cm^2^ (P2) on a TCP culture dish and placed on a humidified incubator (Binder), at 37°C, with 5% CO_2_ – standard culture conditions – or in an InvivO₂® 400 Physoxia Workstation (Baker Ruskinn), with 5% O_2_/5% CO_2_ and humidified environment - physioxia. All procedures were performed within the workstation to avoid oxygen level fluctuations and all reagents used were left to equilibrate for at least 24 hours. When cell culture reached 80-90% confluence, cells were split 1:3 until P4 (Figure 2A).

**Figure 2.**
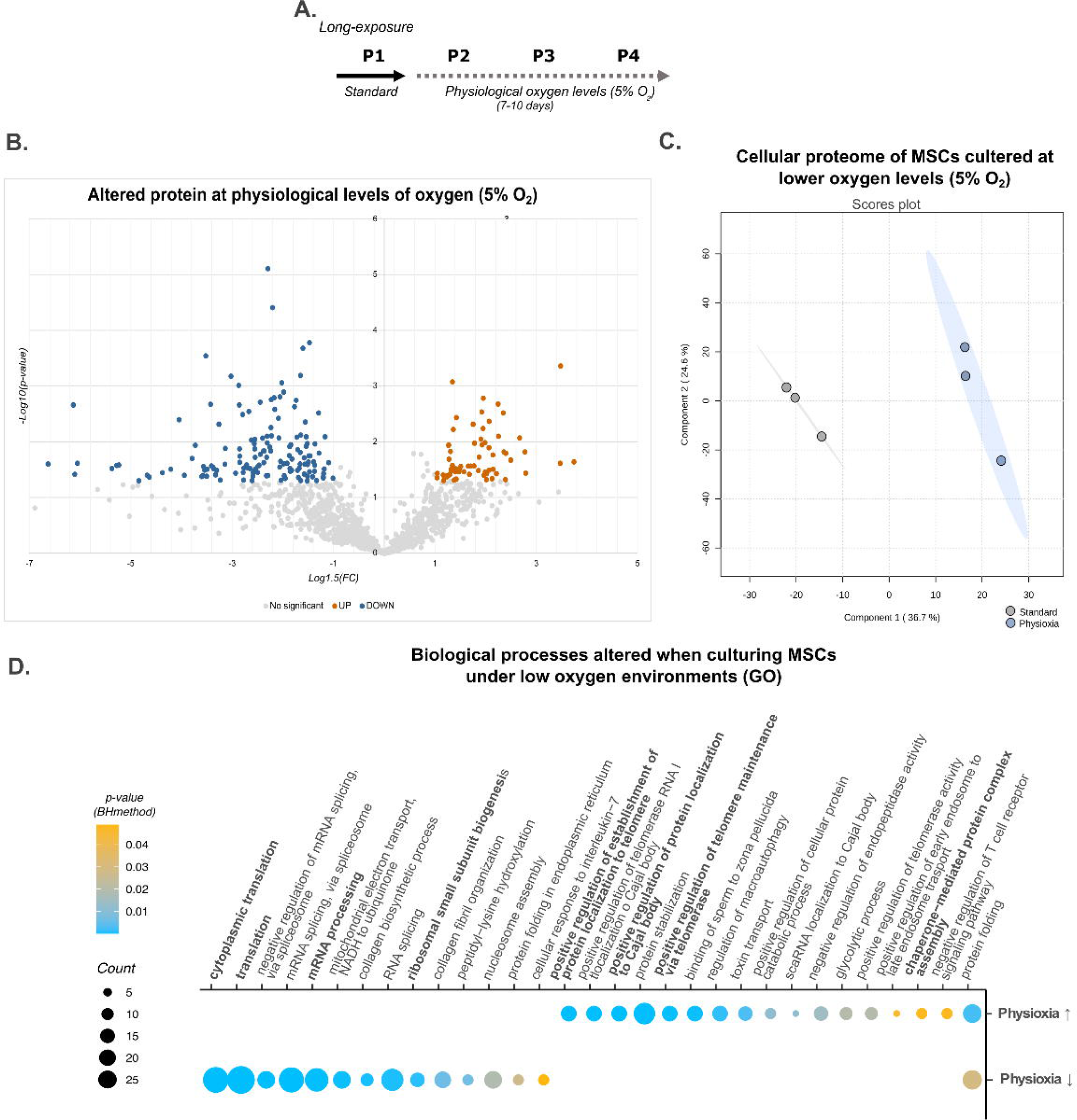
–. MSCs cultured under physiological oxygen levels differentially express proteins belonging to protein homeostasis and translation processes. MSCs were expanded for 7 to 10 days (from P2 to P4) under controlled oxygen levels 5%O_2_), and the cellular proteome was characterized by DDA and DIA LC-MS/MS proteomics (A). The Volcano Plot (B) represents the statistically significant altered proteins (t-test, p<0.05), up-(orange) or down- (blue) regulated (|Log_1.5_FC|>1). Representation of the PLS-DA scores plot (C) for standard (grey) and physioxia (blue) experimental groups using all the quantified proteins. In total, 173 proteins were found to be upregulated, and 310 were downregulated. These proteins were then submitted to a GO analysis (D). The plot illustrates the significant biological processes identified for down- and upregulated proteins, respectively. Abbreviations: Benjamini-Hochberg method (BH).

For priming experiments, UC-MSCs were kept at standard culture conditions until P3 (Figure 4A). Then, cells were seeded at 10,000/cm^2^ (P4) and left at standard conditions or physioxia for 48 hours.

### Proteomic analysis

#### Protein extraction and sample preparation

Protein extracts were collected on ice by the addition of a Lysis buffer (120 mM Tris-HCl pH 6.8, 4% (w/v) Sodium dodecyl sulfate (SDS) and 20% (v/v) glycerol) except for long-exposure mechanomodulated MSCs, where the proteome was fractioned into soluble and membrane fraction (as previously described (24)). Then, samples were denatured and sonicated. The total amount of protein was quantified using Pierce^TM^ 660nm Protein Assay (Thermo Fischer Scientific) according to the manufacturer’s instructions. For in-gel digestion, sample buffer 6× (350mM Tris-HCl pH 6.8, 30% (v/v) glycerol, 10% (w/v) SDS, 0.93% (w/v) dithiothreitol (DTT)) was added to the sample to obtain a final concentration of 1×, and proteins were alkylated by the addition of a 40% acrylamide solution. The same amount of a protein standard MBP-GFP was added to each protein extract (25). For protein identification, pools of each condition were prepared, while for protein quantification, each sample was loaded individually on the gel. Proteins were resolved through a short-GeLC approach and stained with Coomassie Brilliant Blue G-250 (26, 27). Gel lanes were cut into five or three bands, concerning protein identification or quantification, respectively, and destained using a 50 mM ammonium bicarbonate with 30% acetonitrile solution. Proteins were digested overnight, in gel, by trypsin (0.01 mg/ml), and peptides were extracted by the addition of solutions with an increasing percentage of acetonitrile (30, 50 and 98%) and 1% formic acid, dried by vacuum centrifugation at 60°C, and stored at -20°C.

#### Data acquisition

Samples were resuspended on 2% acetonitrile and 0.1% formic acid solution and analyzed on a NanoLC™ 425 System (Eksigent) coupled to a Triple TOF™ 6600 mass spectrometer (Sciex) with an electrospray ionization source (DuoSpray™ Ion Source from Sciex). Peptides were separated on a Triart C18 Capillary Column 1/32” (12 nm, 3 μm, 150 mm × 0.3 mm, YMC, Dinslaken, Germany) and using a Triart C18 Capillary Guard Column (0.5 mm × 5 mm, 3 μm, 12 nm, YMC) at 50 °C. The flow rate was set to 5 µL/min and mobile phases A and B were 5% DMSO plus 0.1% formic acid in water and 5% DMSO plus 0.1% formic acid in acetonitrile, respectively. The LC program was performed as follows: 5 – 30% of B (0 - 50 min), 30 – 98% of B (50 – 52 min), 98% of B (52-54 min), 98 - 5% of B (54 – 56 min), and 5% of B (56 – 65 min). The ionization source was operated in the positive mode set to an ion spray voltage of 5500 V, 25 psi for nebulizer gas 1 (GS1), 10 psi for nebulizer gas 2 (GS2), 25 psi for the curtain gas (CUR), and source temperature (TEM) at 100°C. For Data Dependent Acquisition (DDA) experiments, the mass spectrometer was set to scan full spectra (m/z 350–2250) for 250 ms, followed by up to 100 MS/MS scans (m/z 100–1500) with a 30 ms accumulation time, resulting in a cycle time of 3.3 s. Candidate ions with a charge state between + 1 and + 5 and counts above a minimum threshold of 10 counts per second were isolated for fragmentation, and one MS/MS spectrum was collected before adding those ions to the exclusion list for 15 s (mass spectrometer operated by Analyst® TF 1.8.1, Sciex®). For Data Independent Acquisition (DIA) (or Sequential Window Acquisition of all Theoretical Mass Spectra (SWATH)) experiments, the mass spectrometer was operated in a looped product ion mode and specifically tuned to a set of 42 overlapping windows, covering the precursor mass range of 350–1400 m/z. A 50 ms survey scan (m/z 350–2250) was acquired at the beginning of each cycle, and SWATH-MS/MS spectra were collected from 100 to 2250 m/z for 50 ms, resulting in a cycle time of 2.2 s.

#### Data analysis

Each fraction of pooled samples was acquired individually in DDA mode. ProteinPilot™ software (v5.0, Sciex, Framingham, MA, USA) was used to generate the ion library of the precursor masses and fragment ions by combining all files from the pools. The data was searched against the reviewed Human (Swiss-Prot) database (downloaded on 21^st^ October 2021 (long-exposure experiments) and 16^th^ May 2022 (priming)). The quality of the identifications was assessed by an independent False Discovery Rate (FDR) analysis performed using the target-decoy approach provided by the software.

Samples were acquired on DIA mode (SWATH) for protein relative quantification. Data was processed using the SWATH™ plug-in for PeakView™ (v2.0.01, Sciex®), setting 15 peptides per protein and five transitions per peptide. Only peptides present in at least one-third of the samples per condition with an FDR below 1% were quantified. Data was normalized to the total intensity. The MS data have been deposited to the ProteomeXchange Consortium via the PRIDE partner repository with the dataset identifier PXD047973 and PXD017674 (28).

Statistical analysis was performed on MetaboAnalyst 5.0 (29). Proteins of interest were selected through multivariate (Partial least squares-discriminant analysis (PLS-DA)) or univariate (t-test or Mann-Whitney or Kruskal-Wallis tests) analysis. Proteins of interest were selected if they had a VIP>1 or p<0.05 and fold change (FC) above 1.5 or below 0.67 (|Log_1.5_FC|>1) and subjected to a Gene Ontology enrichment analysis on Funrich (30). Reactome database was used to map pathways (31), Venn diagrams were generated on DeepVenn (32), and violin plots were created on GraphPad Prism 9.

## Results

### Physiological stiffness and oxygen levels alter common proteins involved in translation

Mimicking physiological stiffness and oxygen levels *in vitro* impacts cell proteome. To understand the impact of each, first, each cue was analyzed separately. The data concerning the mechanomodulated MSCs (24) was reanalyzed for the purpose of the present study. First, the proteome of UC-MSCs expanded from P2 to P4 on a soft substrate (3kPa) or standard stiff culture conditions (GPa) was characterized (Figure 1A). Of the 1452 identified proteins, 698 and 763 were quantified with confidence in membrane and soluble fractions, respectively. PLS-DA models were generated for each fraction without overlapping standard and softer stiffness conditions (Figure 1B, C). Then, proteins of interest selected for each fraction were compared, denoting that few proteins were quantified on both fractions (Figure 1D). Filtered proteins were then subjected to GO analysis (Figure 1E). Briefly, proteins associated with cell redox homeostasis, such as peroxiredoxins or thioredoxins, including specific subtypes found in mitochondria, were upregulated (Supplementary Table 1). In contrast, proteins involved in the proteossome, translation and ribosomal subunit biogenesis processes were found to be downregulated. Particularly, many proteins associated with these GOs are ribosomal subunits (Supplementary Table 1). Together, these data suggest that culturing UC-MSCs on low stiffness (3 kPa) can profoundly modulate several metabolic processes, including redox and protein homeostasis.

To explore the impact of oxygen levels on UC-MSCs *in vitro* expansion, the cellular proteome of UC-MSCs cultured under controlled levels of oxygen (∼5%O_2_) and standard culture conditions (∼18%O_2_) (P2 to P4, for 7 to 10 days) was analyzed (Figure 2A, Supplementary Figure 1A). A total of 3107 proteins were identified, from which 1234 were quantified with confidence. Both univariate and multivariate analyses (Figure 2B, C) revealed a differential expression of several proteins’ levels after exposure to more physiological oxygen levels, showing that physioxia can modulate the cellular proteome. To further investigate which processes were being altered, proteins that were statistically relevant and more or less abundant (|Log_1.5_FC|>1) were subjected to GO enrichment analysis (Figure 2D). Interestingly, physioxia leads to increased levels of proteins associated to protein catabolic processes and telomerase maintenance-associated proteins, including several chaperones, such as Heat shock protein HSP90 and T-complex proteins (Supplementary Table 2). In contrast, the levels of proteins involved in RNA splicing appear to be downregulated. These results indicate that if lowering the oxygen levels is interfering with both RNA and protein metabolism, possibly *in vitro* standard culture conditions might be dysregulating these mechanisms. Moreover, translation and ribosomal biogenesis processes were also downregulated, presenting the same tendency described on MSCs cultured on compliant stiffness platforms (Figure 1E). Based on these common findings encountered on these two distinct physiological stimuli, the proteomes on each experimental setup were compared to explore other possible matches.

Reducing the stiffness or oxygen levels in the culture conditions to values more closely aligned with physiological conditions results in a consistent downregulation of proteins associated to translation and ribosomal biogenesis. To deeply explore if these physiological cues would culminate on the modulation of the same processes, the proteins of interest previously identified in each experimental setup were compar0ed (Figure 3A). Then, each of the matches was subjected to a GO enrichment analysis (except for physioxia downregulated proteins and mechanomodulated upregulated proteins, where no statistically significant enriched GO hits were found). Most of the shared proteins were down-regulated both on mechanomodulated or physioxia-cultured MSCs, and, not surprisingly, were associated with ribosomes and translation processes (Figure 3B). Although the number of common upregulated proteins was small, it was still possible to confirm that they were mainly related to mitochondrial ATP synthesis processes, suggesting that exposing MSCs to standard culture conditions might also be affecting the energy pathways (Figure 3C). In contrast, some proteins related to the actin cytoskeleton and focal adhesions displayed inverse tendencies on both culture conditions (Figure 3D). This piece of evidence indicates a crosstalk between oxygen-sensor pathways and MSCs cytoskeleton, a topic that is gaining interest and is known to interfere with the differentiation mechanisms of stem cells (33, 34).

**Figure 3.**
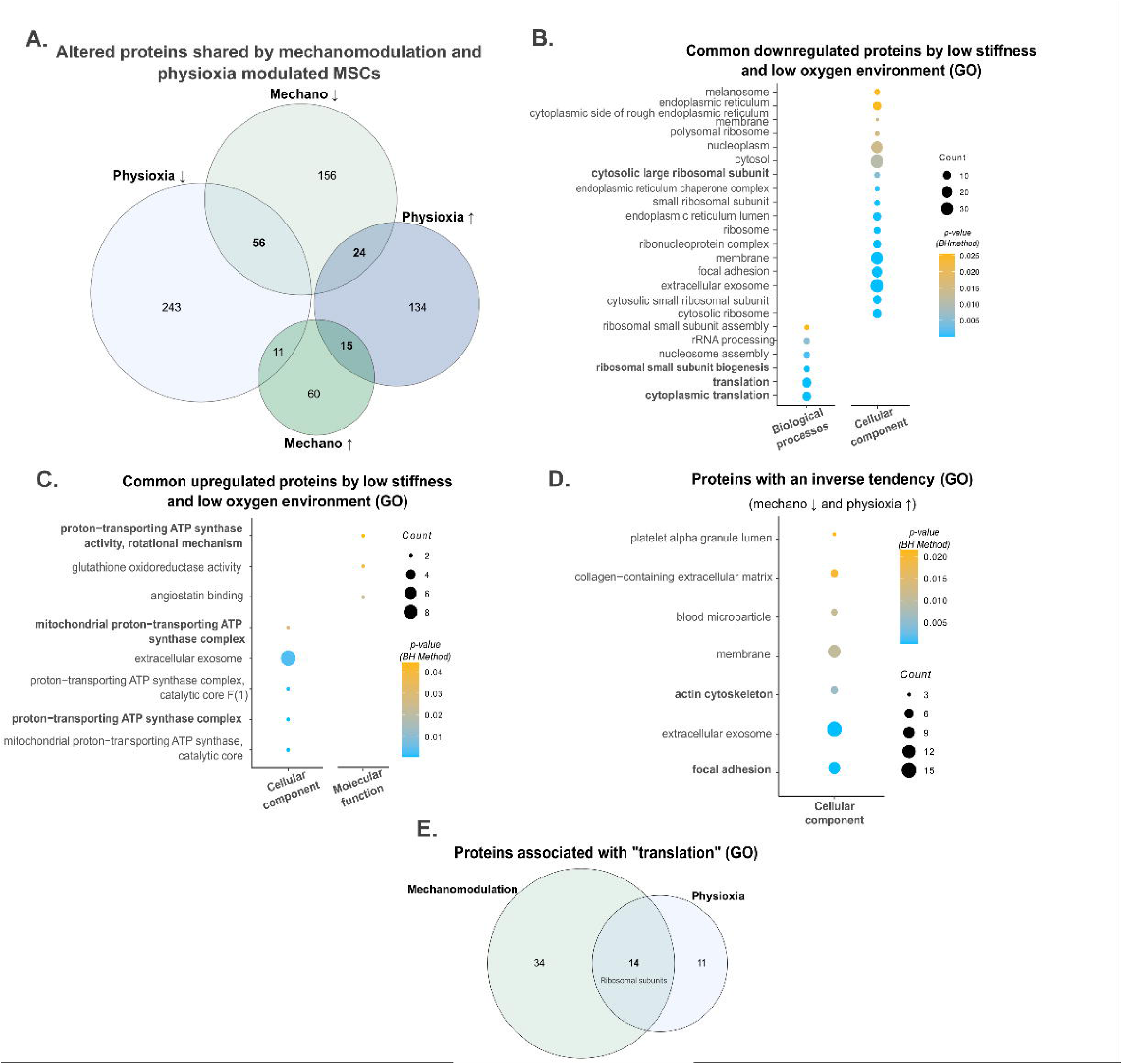
–. *Oxygen levels (5%O_2_) and stiffness (3kPa) closer to physiological levels trigger common pathways on MSCs.* (A) Selected proteins of interest (|Log_1.5_FC|>1) on each individual analysis and when combined. Mechanomodulation (Mechano) is represented in green, while physioxia is depicted in blue. To identify common significant biological processes and structures altered, GO analysis was performed using the shared proteins modulated by these different cues (B, C, D). Data analysis revealed that the shared proteins associated with “translation” were fourteen ribosomal subunits (Supplementary Table 3) (E). Abbreviations: Benjamini-Hochberg method (BH); Mechanomodulation (Mechano).

Finally, since a downregulation of translation triggered by lower stiffness and oxygen levels was highlighted in the previous analysis, the proteins associated with this GO hit were compared (Figure 3E). In fact, fourteen proteins from this ontology were commonly altered in these distinct physiological conditions when compared to the conventional cell culture condition and are 60S and 40S ribosomal subunits (Supplementary Table 3). Together, these data show that despite being two different *in vitro* environments mimicking distinct physiological conditions, they converge on the modulation of ribosomal subunit expression, implying that expanding these cells under standard culture conditions is interfering with MSCs’ protein metabolism.

### UC-MSCs respond more promptly to low stiffness environments, while they need more time to adapt to low oxygen levels

Data presented in the previous section described the cellular proteome alterations on naïve UC-MSCs cultured permanently under more physiological culture conditions, either stiffness or oxygen levels (7 to 10 days (P2-P4)). However, it has been described that MSCs display mechanical memory if cultured for long periods (>10 days) on a stiff substrate (35). Therefore, to clarify if these cells still had the ability to respond to lower stiffness and oxygen levels after being expanded under standard culture conditions, UC-MSCs at P4 were moved to lower stiffness or oxygen levels environments for 48 hours (priming), since UC-MSCs are known to adapt to the environment stiffness on the first 24 hours (36) (Figure 4A). The proteome was then collected and analyzed by LC-MS/MS proteomics workflow to (i) identify initial mechanisms triggered by these cues, (ii) accommodate experimental and donors’ variability by the use of matching donors for the same experimental conditions, (iii) optimize an easier and economically viable protocol to modulate UC-MSCs. Mimicking physiological conditions *in vitro* requires specialized equipment, and it is time-consuming since the proliferation rate of these cells is low (Supplementary Figure 1B,C). Out of the 1992 proteins identified, nearly 1300 proteins were quantified with confidence. A Principal Component Analysis (PCA) revealed a separation between the mechanomodulation group and the other conditions, physioxia and standard, which overlap (Figure 4B). The results are identical when using a clustering analysis (Supplementary Figure 2A), which displays a closer relationship between physioxia and standard culture conditions. For both cues, the total number of downregulated proteins was higher than upregulated proteins (Figure 4C, Supplementary Table 4). Although a limited number of proteins was initially shared between conditions, the common processes altered upon long exposure, as translation, suggest that different players will converge on the modulation of the same mechanisms. The GO enrichment analysis revealed that altered proteins upon priming were primarily associated to extracellular components, particularly when responding to lower stiffness environments (Figure 4D). Accordingly, downregulated proteins also indicate rearrangements in the extracellular matrix organization and cell adhesion mechanisms (Figure 4D). Taken together, analysis of the primed UC-MSCs proteome demonstrates that mechanomodulation is a more vigorous stimulus that can modulate the interaction of UC-MSCs with the extracellular environment and some cellular compartments, as the endoplasmic reticulum, within the first 48 hours. The closer relationship on the proteins’ expression profile of UC-MSCs cultured under lower oxygen levels, compared to standard conditions, as well as the reduced number of altered proteins (62 out of 1300 quantified) and the low number of significant GO hits indicate that UC-MSCs require prolonged exposure periods to adapt their cellular machinery to this culture conditions. Notwithstanding, by generating a PLS-DA model for each condition (Supplementary Figure 2B and C), it was still possible to distinguish cells cultured under standard conditions from cells exposed to lower stiffness or lower oxygen levels.

**Figure 4.**
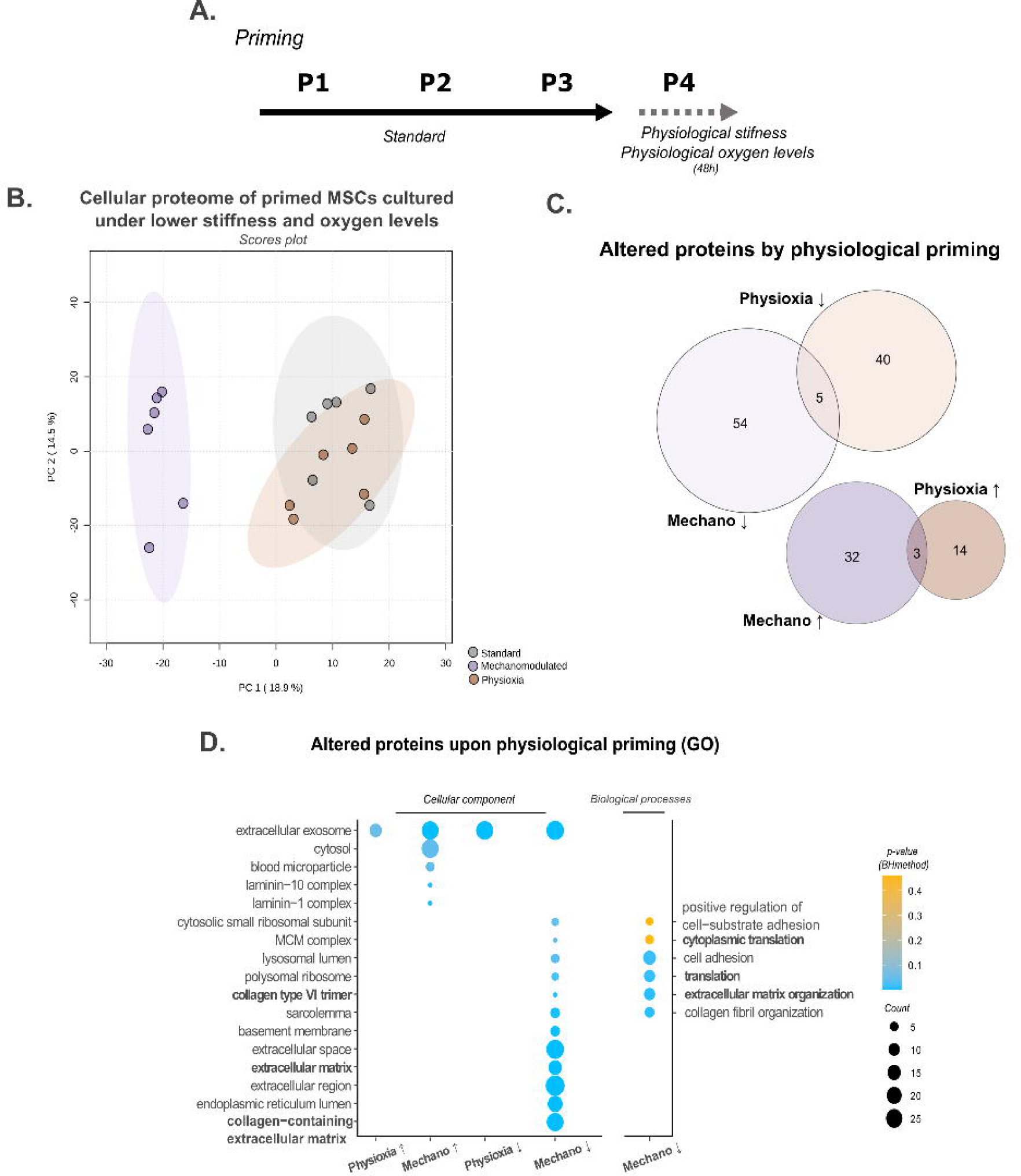
–. *UC-MSCs can readjust their cellular proteome after being expanded under standard culture conditions.* UC-MSCs were expanded under standard culture conditions and moved to lower stiffness (3kPa) or oxygen levels (5%O₂) culture conditions for 48h – priming (A) at P4. All proteins quantified with confidence in the three experimental groups (standard (grey), physioxia (brown) and mechanomodulation (purple)) are represented on the PCA scores plot (B). Proteins of interest (|Log_1.5_FC|>1) were compared (C) and subjected to a GO enrichment analysis. Significant cellular components and biological processes are represented (D). Abbreviations: Benjamini-Hochberg method (BH); Mechanomodulation (Mechano).

To investigate whether the initial expression patterns were kept when UC-MSCs were cultured under an environment that mimics physiological stiffness or oxygen levels, relevant proteins altered by the priming and long exposure were compared. As expected, cells cultured in a more physiological environment for extended durations exhibit a greater overall number of modified proteins (Figure 5A and B; Supplementary Tables 5 and 6), suggesting a deeper cellular rearrangement. While only 11(3.87% of the total) and 7 (2.01% of the total) were commonly downregulated upon mechanomodulation and physioxia, respectively, they might provide insights into the initial response and adaptation to these culture conditions. The highlighted proteins are involved in the relevant biological processes identified in long exposure experiments, such as cell adhesion, translation, and cell redox homeostasis (Figure 5C, D; Supplementary Table 1,2). Versican Core Protein (P13611) and Collagen alpha-2 (VI) chain (P12110) are extracellular matrix proteins and are downregulated at both short and long-term points in the lower stiffness conditions, confirming that matrix components modifications are crucial to the adaptation of UC-MSCs to mechanical cues of the environment. Also, 60s ribosomal protein L27 (P61353), a modulated protein involved in translation processes, responded to low oxygen levels of the culture condition in the first 48h and long exposure. Finally, for mechanomodulated UC-MSCs, thioredoxin (P10599) was found to be upregulated on UC-MSCs cultured on a soft substrate at both time points but unchanged under both physioxia conditions. A better understanding of the role of these proteins in transducing physiological cues could provide insightful knowledge to empower the therapeutic potential of UC-MSCs.

**Figure 5.**
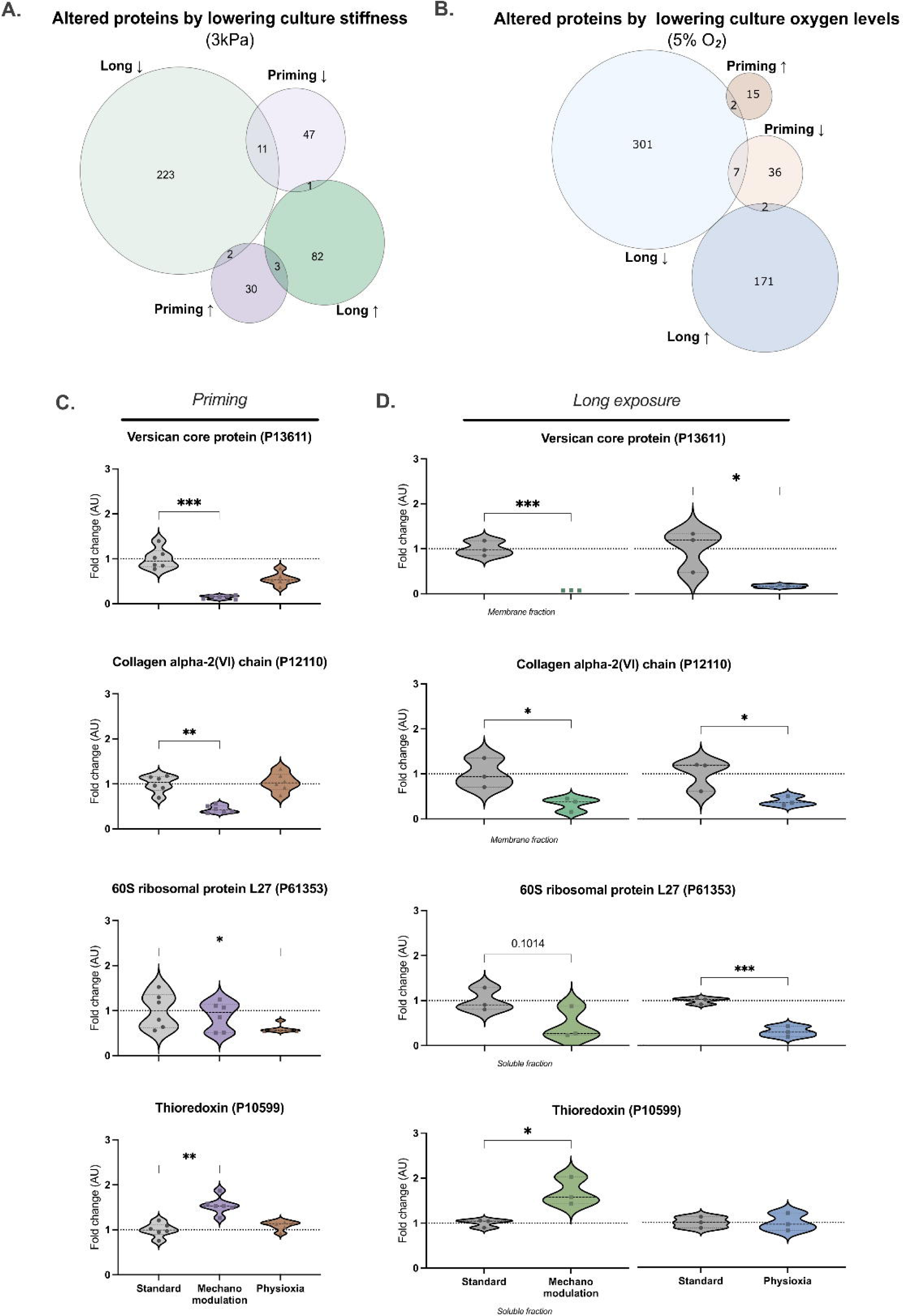
*Extracellular matrix organization and translation proteins maintain their expression pattern consistent with prolonged exposure.* For both stimuli, only a small number of proteins were kept significantly altered on priming and long exposure to physiological stimuli (Figure 5A, B). Violin plots highlight proteins that belong to the extracellular matrix, such as Versican core protein (C) or Collagen alpha-2(VI) chain (D), and translation, as 60S ribosomal protein L27 (E). The increased antioxidant thioredoxin expression is also kept in priming and long-exposure experiments (F). All highlighted proteins were previously selected by statistical tests (p<0.05 and/or VIP>1), except for Thioredoxin after long-exposure to physioxia. (Kruskal-Wallis and t-test were applied on priming and long-exposure experiments, respectively; *p<0.05, **p<0.01)

### Lower stiffness or oxygen levels modulate proteins involved in cellular senescence pathways

The proteomic characterization of UC-MSCs maintained at lower oxygen levels revealed that proteins involved in telomerase maintenance were upregulated (Figure 2D). The shortening of telomeres is known to be associated with cell senescence or permanent cell cycle arrest (37). To further explore if mimicking physiological culture conditions could have an impact on cellular senescent phenotype, the Reactome database was used to map this pathway (R-HSA-2559583) (Supplementary Table 8). Although the pathway was not significantly altered, mapped protein IDs were extracted and analyzed individually. Priming UC-MSCs on soft substrates resulted in an increase of transcription factor p65 (RELA) (part of NF-κB signaling pathway) and high mobility protein HMG-I/Y, suggesting modifications at the transcriptional level (Figure 6A, B and S2A-C). Interestingly, for both long and short exposure periods to low stiffness or oxygen levels, specific histone subtypes also appear downregulated (Figure 6C, D). Therefore, it is possible to infer that apart from modification on the redox homeostasis and protein metabolism, culture conditions mimicking physiological conditions also interfere with DNA organization. Taken together, the results suggest that lowering the stiffness or oxygen levels of culture conditions leads to a cellular rearrangement of several cellular processes that together might impact MSCs aging during *in vitro* expansion (Figure 6E).

**Figure 6.**
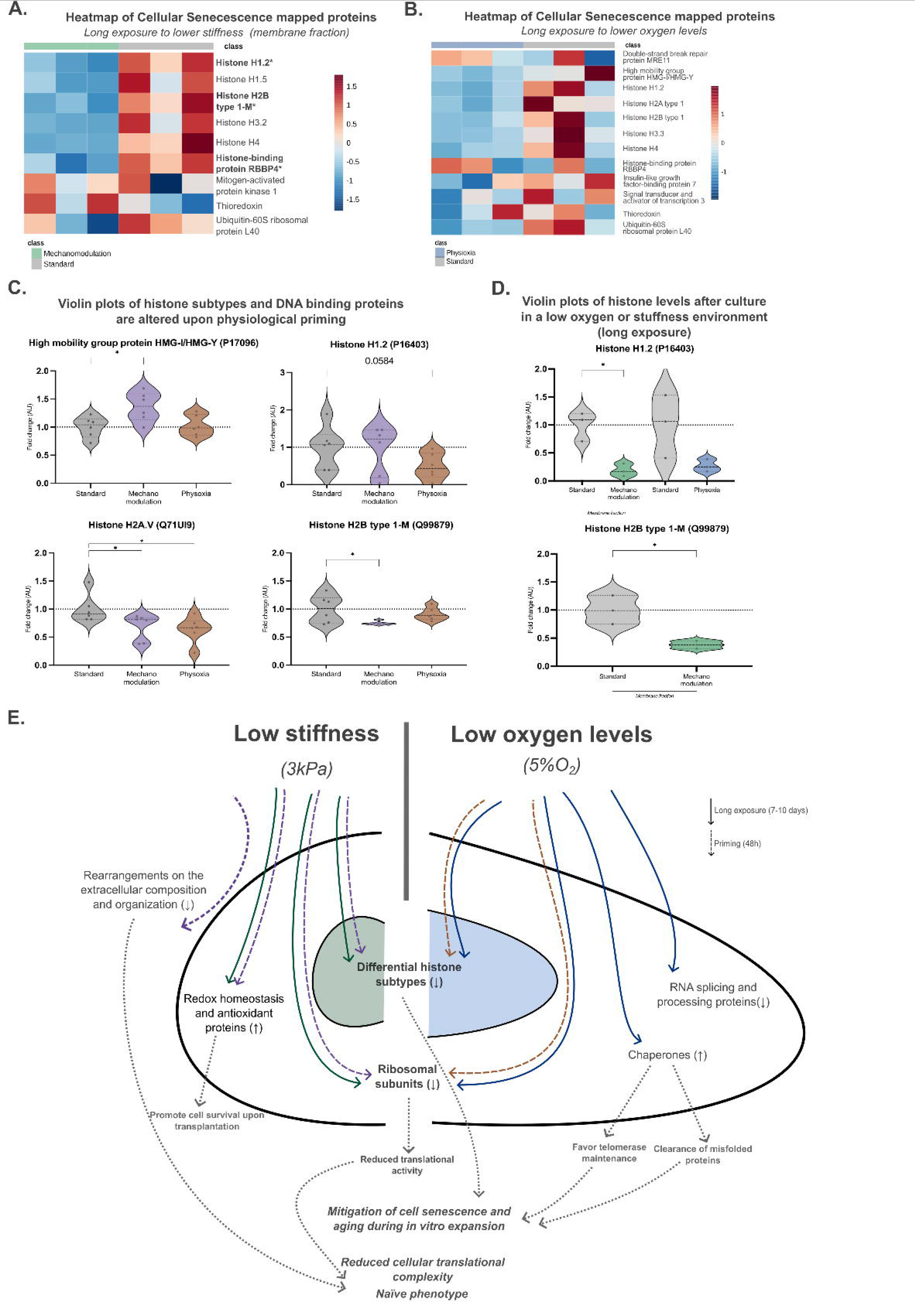
*Lower stiffness and oxygen levels reorganize UC-MSCs’ machinery associated with cell senescence*. Highlighted proteins from previous results were analyzed on Reactome. Mapped protein IDs associated with Cellular senescence (R-HSA-2559583) were analyzed individually. Heatmaps plot the mapped protein levels on each screening (normalized to control) of UC-MSCs cultured at lower stiffness and oxygen levels for long periods (A, B, respectively). Violin plots highlight significantly altered protein levels upon priming (C, D) and long exposure (E). Kruskal-Wallis and t-tests were applied on priming and long-exposure experiments, respectively; *p<0.05. Schematic representation of the mechanisms potentially regulated by long (7-10 days) or short (48h) exposure to lower stiffness and oxygen levels during cell expansion (I). Arrows with continuous line (→) - proteins altered by long exposure; Arrows with dashed lines (→) proteins altered by priming; Arrows with doted lines (→)– hypothetical implications on cellular processes, according to the literature).

## Discussion

The clinical use of MSCs holds great promise as a therapeutic strategy for various diseases (3). However, producing a single treatment dose can be challenging due to the expensive and time-consuming resources required. Additionally, extensive *in vitro* expansion may compromise their properties (10). To address these challenges, it was hypothesized that mimicking the physiological environment (3kPa or 5%O_2_, respectively) could enhance the therapeutic potential of MSCs since it is described that lowering the stiffness and oxygen levels promote MSCs stemness (36, 38). To explore how these culture conditions would impact these cells, the cellular proteome of physiologically modulated UC-MSCs was characterized.

The first step was to characterize the proteome alterations resulting from a long-term culture of the cells under physiological conditions (P2 to P4; 7 to 10 days). Proteins related to cell redox homeostasis were upregulated by a softer environment (Figure 1D), including the thioredoxin antioxidant system (Supplementary Table 1). These proteins act as scavengers of reactive oxygen species (ROS) and play a crucial role in mediating cellular response to oxidative stress response (39, 40). Its levels were altered in response to long and short stimuli, indicating its involvement in the initial pathways triggered by stiffness (Figure 5C, Supplementary Table 1 and 5). When MSCs are transplanted, they often encounter a harsh environment at the injury site, resulting in the loss of more than 90% of cells within the first 24 hours, with oxidative stress the major contributor (41). Therefore, an increased expression of redox proteins might play a crucial role in MSCs’ survival. More precisely, Suresh *et* al. demonstrated promising results on a rat model of infarcted myocardium using Thioredoxin-1 engineered MSC, which were found to reduce fibrosis and increase the release of pro-angiogenic factors (42). Similar positive outcomes were observed in the context of scleroderma, when using bone marrow MSCs expressing Thioredoxin-1 (43). An advantage of the mechanomodulation of MSCs is the simultaneous modulation of proteins of the redox machinery, including peroxiredoxins or thioredoxin reductases. Therefore, it is possible to infer that mechanomodulated MSCs might be more resilient to harsh conditions post-transplantation (44, 45).

Moreover, controlling oxygen levels in culture has become a prominent subject in scientific research, especially concerning their impact on MSCs (46, 47). To our knowledge, the proteomic characterization, elucidating cellular alterations induced by physiological oxygen levels *in vitro*, has not been addressed thus far. Therefore, it is particularly interesting that the data presented in this study indicate that the proteome is deeply altered when MSCs are cultured at 5% O_2_, especially for long periods (P2 to P4, 7-10 days). More specifically, subunits of the chaperonin-containing T-complex (TRiC) were found to be upregulated alongside other chaperones, such as HSP90 (Figure 2D, Supplementary Table 2). Chaperones play a crucial role in maintaining proteostasis and are associated with cellular aging, since senescent MSCs have been described to exhibit dysregulation in the chaperone network, leading to the accumulation of misfolded proteins (48, 49). Also, late passage MSCs (P5-P18) are known to express reduced levels of both TRiC and HSP90 (49). In addition, these chaperones are also described to be implicated in telomere maintenance, a finding supported by our GO enrichment data (Figure 2C) (50, 51). Despite the low proliferation rate of UC-MSCs kept in low oxygen environments, they are not positive for β-galactosidase, which is in line with a shift to a quiescent phenotype (Supplementary Figure 1D) (52). While complementary approaches are required to confirm telomere integrity and the clearance capacity of misfolded proteins after MSCs expansion under low oxygen levels, these data unveil innovative strategies to uphold protein homeostasis and preserve telomere integrity, potentially mitigating cell senescence and aging while being expanded *in vitro*.

Stiffness and oxygen levels are two distinct stimuli, yet the proteomic comparison here presented revealed a shared regulatory process influenced by these physiological cues: the downregulation of several ribosomal subunits and consequently influencing translation processes (Figure 3C). Not only 60S ribosomal protein L27 (P61353) but other ribosomal proteins critical to rRNA processing are downregulated at priming and long culture conditions under both mechanomodulation and physiological oxygen levels (Figures 4D and 5E) (53). It has been described that ribosome concentration can favor the expression of specific mRNA subtypes and, consequently, protein expression (ribosome concentration hypothesis) (54, 55). Accordingly, it is now established that translation plays a significant role in determining stem cell fate (55). After initiating differentiation, embryonic stem cells increase their cellular complexity in terms of endoplasmic reticulum and Golgi complex and exhibit a higher translation activity rate, regulated via mTOR signaling pathway (56, 57). Consequently, replicating physiological stiffness and oxygen levels *in vitro* could potentially promote a more naïve phenotype on MSCs through the downregulation of ribosomal subunit expression. However, the translation activity rate still needs to be validated by complementary techniques.

UC-MSCs at low passage (until P4) can adapt and expand under physiological conditions *in vitro*. However, whether UC-MSCs could still respond to these cues and what initial adaptation mechanisms might be modulated after standard conditions expansion (2-3GPa, 18%O_2_) were questioned. Stiffness appears to induce a more profound modulation of the proteome of UC-MSCs within the first 48 hours, implying that these cells need more time to fully adapt to physiological oxygen, as it was proposed for human umbilical vein (HUVEC) cells, which require at least five days to fully adapt to the environment (Figure 4B) (58). However, despite the increased levels of YAP transcription factor, known to be mechanosensitive, CTGF (a downstream target of YAP) and cofilin/phosphorylated-cofilin ratio remain unaltered on both experimental conditions, reinforcing that long periods are necessary for UC-MSCs fully adapt their intracellular machinery (35) (Supplementary Figure 2D,E,F). Nevertheless, during this short timeframe, GO analysis shows a rearrangement at the extracellular matrix level in mechanomodulated primed MSCs (Figure 4D). In detail, both Versican Core Protein (P13611) and Collagen alpha-2(VI) chain (P12110) levels were decreased under more physiological conditions during priming and long-term culture (Figure 5C, D). The increased levels of these proteins on the extracellular matrix are described to favor a commitment to chondrogenic differentiation (59, 60). Also, the collagen deposition leads to an increase on matrix stiffness, impacting MSCs differentiation potential (61). Therefore, it can be inferred that extracellular matrix components play a pivotal role in MSCs’ adaptation to physiological culture conditions.

All these findings lead to a more in-depth investigation into proteins associated with cell senescence. The NF-_κ_B pathway, through p65 transcription factor, is documented in the literature as being responsive to microenvironment oxygen levels and playing a role in replicative senescence and differentiation (62-64). Since it appears to be involved in MSCs’ adaptation to lower stiffness and oxygen levels, it might be an interesting target to investigate. Furthermore, reducing stiffness leads to an increased expression of the high mobility group protein HMG-I/Y, which is known to promote the expression of pluripotency markers (65). Additionally, chromatin modifications are manifested through the downregulation of various histone subtypes in MSCs cultured for both long and short periods. These proteins are responsible for packing genomic DNA in the nucleus, and their functions are highly dependent on post-translational modifications (66). More specifically, the accumulation of histone variants H2A and H2B was described to be associated with a senescent phenotype (67, 68). Thus, exploring the patterns of histone modifications cultured under conditions that mimic physiological stiffness and oxygen levels could yield valuable insights into how standard culture conditions might bias MSCs behavior *in vitro* and how it can be reverted to prevent cellular senescence and/or maintenance of a stemness phenotype.

Taken together, data presented in this study suggests that expanding UC-MSCs under conventional culture conditions (2-3GPa, 18.5% O_2_) is affecting several cellular processes. By contrast, the prolonged expansion of UC-MSCs under low stiffness and oxygen levels environment seems to promote a more naïve phenotype, characterized by a differential expression on histones’ profile and a reduction in the expression of ribosomal subunits (Figure 6E). However, each stimulus has its unique effects: mechanical cues modulate cell redox homeostasis, which might support cell survival at the site of injury, while the reduction of the oxygen levels appears to counteract aging and senescence by altering protein homeostasis (69). Future studies should complement how these metabolic shifts may enhance the therapeutic potential of MSCs and influence the expression of differentiation capacity and stemness markers.

Adopting culture conditions that mimic the physiological environment of UC-MSCs could serve as an alternative, ensuring the preservation of the therapeutic potential of these cells during in vitro expansion. Nevertheless, it should be taken into consideration that recreating their physiological environment is expensive, time-consuming, and requires specialized equipment. The synergistic combination of mechamodulation and physioxia could offer an alternative, particularly when considering priming. Additionally, investigating how these physiological cues influence the secretion of proteins can yield further insights into the paracrine effects triggered by MSCs.

## Supporting information

Methods in detail

## List of abbreviations

BH: Benjamini-Hochberg method
COL-I: Type-I collagen
DDA: Data Dependent Acquisition
DIA: Data Independent Acquisition
DTT: Dithiothreitol
E: Young’s modulus
FBS: Fetal bovine serum
FC: Fold change
FDR: False Discovery Rate
FN: Fibronectin
HPL: Human platelet lysate
HUVEC: Human umbilical vein
LC-MS/MS: Liquid chromatography coupled to tandem mass spectrometry
MEM-α: Minimum essential Medium-α
MSCs: Mesenchymal stem cells
PBS: Phosphate Buffered Saline
PCA: Principal component analysis
PDMS: Polydimethylsiloxane
PLS-DA: Partial least squares-discriminant analysis
UC-MSCs: Umbilical cord mesenchymal stem cells
ROS: Reactive oxygen species
SDS: Sodium dodecyl sulfate
SWATH: Sequential Window Acquisition of all Theoretical Mass Spectra
TCP: Tissue-culture polystyrene
TRiC: Chaperonin-containing T-complex
UC: Umbilical cord

## Funding

This work was financed via FCT – Fundação para a Ciência e a Tecnologia, under projects POCI-01-0145-FEDER-029311, POCI-01-0247-FEDER-045311, UIDB/04539/2020, UIDP/04539/2020, and 10.54499/EXPL/BTM-TEC/1407/2021. SIA was supported by the CEEC grant 10.54499/2021.04378.CEECIND/CP1656/CT0011. IC and CD were supported by individual Ph.D. fellowships, 10.54499/SFRH/BD/143442/2019 and SFRH/BD/115527/2016, respectively. Also, this project was supported by the “la Caixa” Foundation - within the scope of the Promove grant, held in collaboration with BPI and partnership with the Foundation for Science and Technology, REPAIR - PL23-00001.

## Declaration of generative AI and AI-assisted technologies in the writing process

During the preparation of this work the authors used ChatGPT in order to check spelling and correct grammar errors. After using this tool/service, the authors reviewed and edited the content as needed and take full responsibility for the content of the publication.

---

**Supplementary Figure 1.**
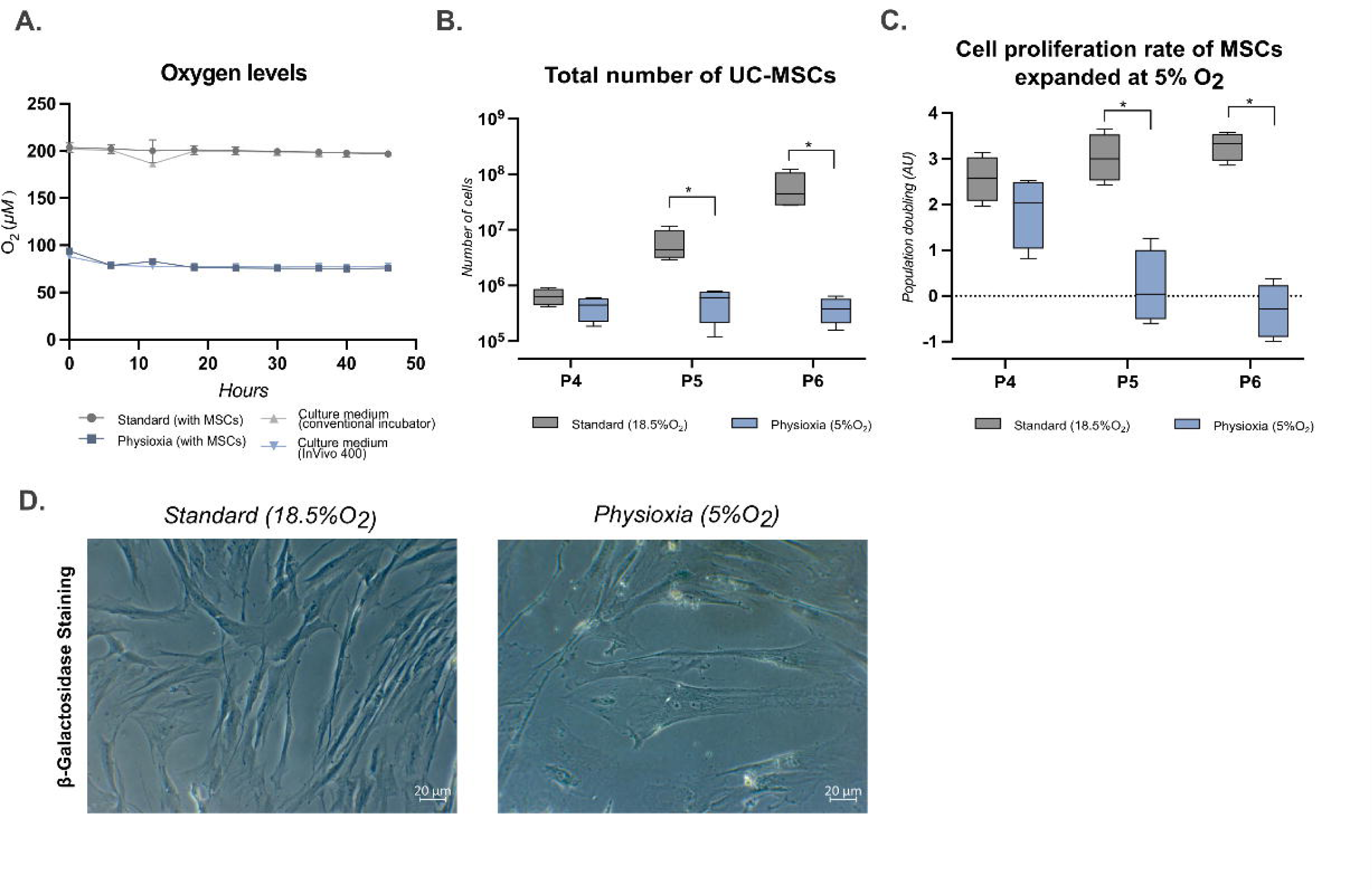
*Proliferation kinetics and senescence assay of UC-MSCs cultured under 5% O_2_.* The oxygen level present in the culture media was, on average, 78.80 µM (5.81% O_2_) for physioxia (blue) and 198.03 µM (14.65% O_2_) (grey) for standard culture conditions (normoxia) (A). The total number of cells was calculated and the population doubling of MSCs expanded at lower oxygen levels (B, C) (Mann-Whitney, * p<0.05). Both experimental conditions were stained for β-galactosidase to assess cell senescence (D).

**Supplementary Figure 2.**
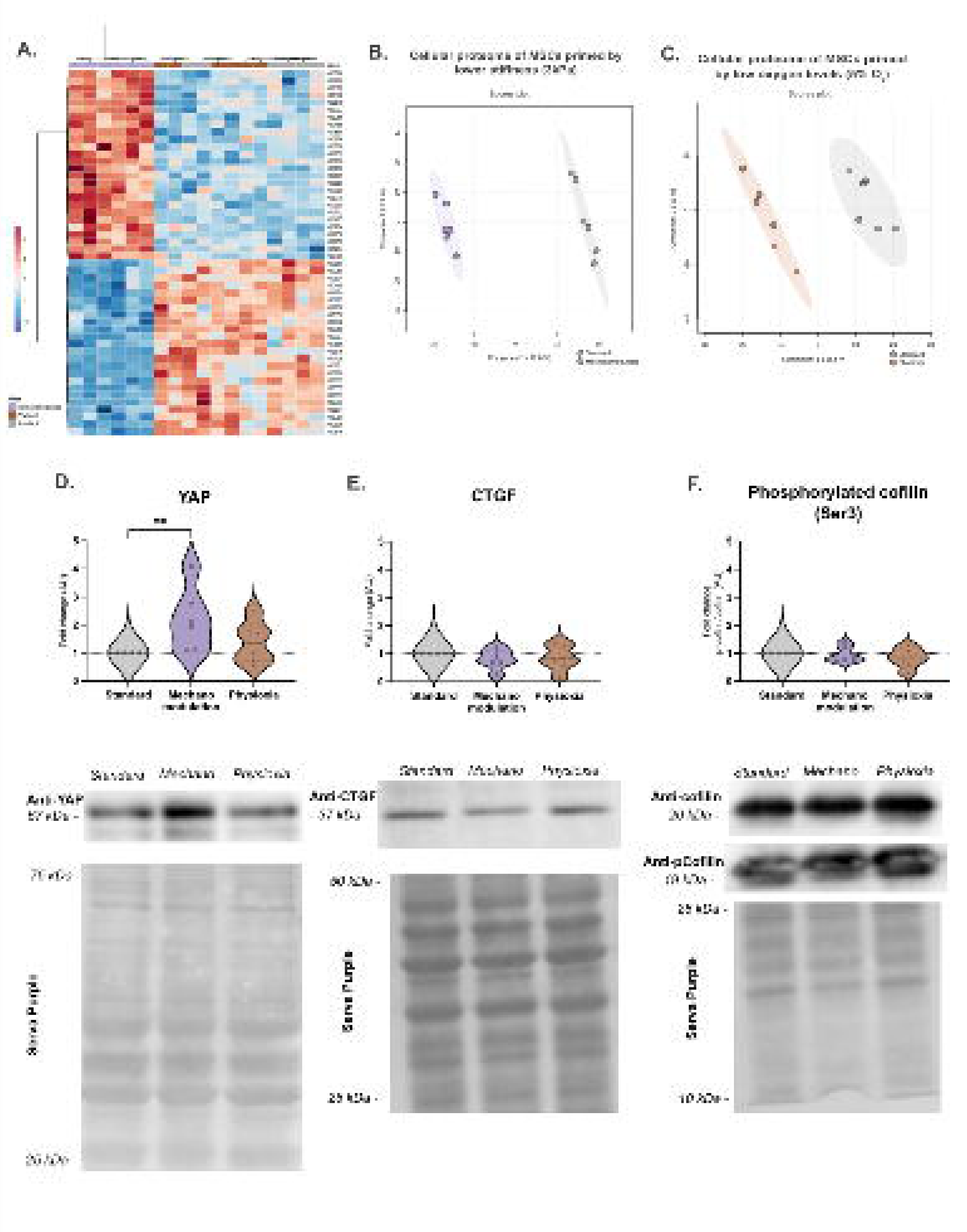
*Priming UC-MSCs with physiological environment (3kPa or 5%O_2_) alters the proteomic profile.* The heatmap represents the TOP 50 proteins (VIP>1) (A). All proteins quantified with confidence were used to generate two distinct PLS-DA models for mechanomodulation (B) and physioxia (C) experimental conditions. Standard culture conditions are represented in grey, while mechanomodulation and physioxia are represented in purple and brown, respectively. Proteins involved in mechanotransduction signaling (YAP (D), CTGF (E), cofilin and phosphorylated-cofilin (Ser3) (F)) were quantified by Western Blot. Protein relative quantification is represented on the violin plots (upper panel) (Kruskall-Wallis test, ** p<0.01). Middle and lower panels display representative images of the bands and total protein staining (Serva purple), respectively. (Abbreviations: Mechanomodolation (Mechano).

**Supplementary Figure 3.**
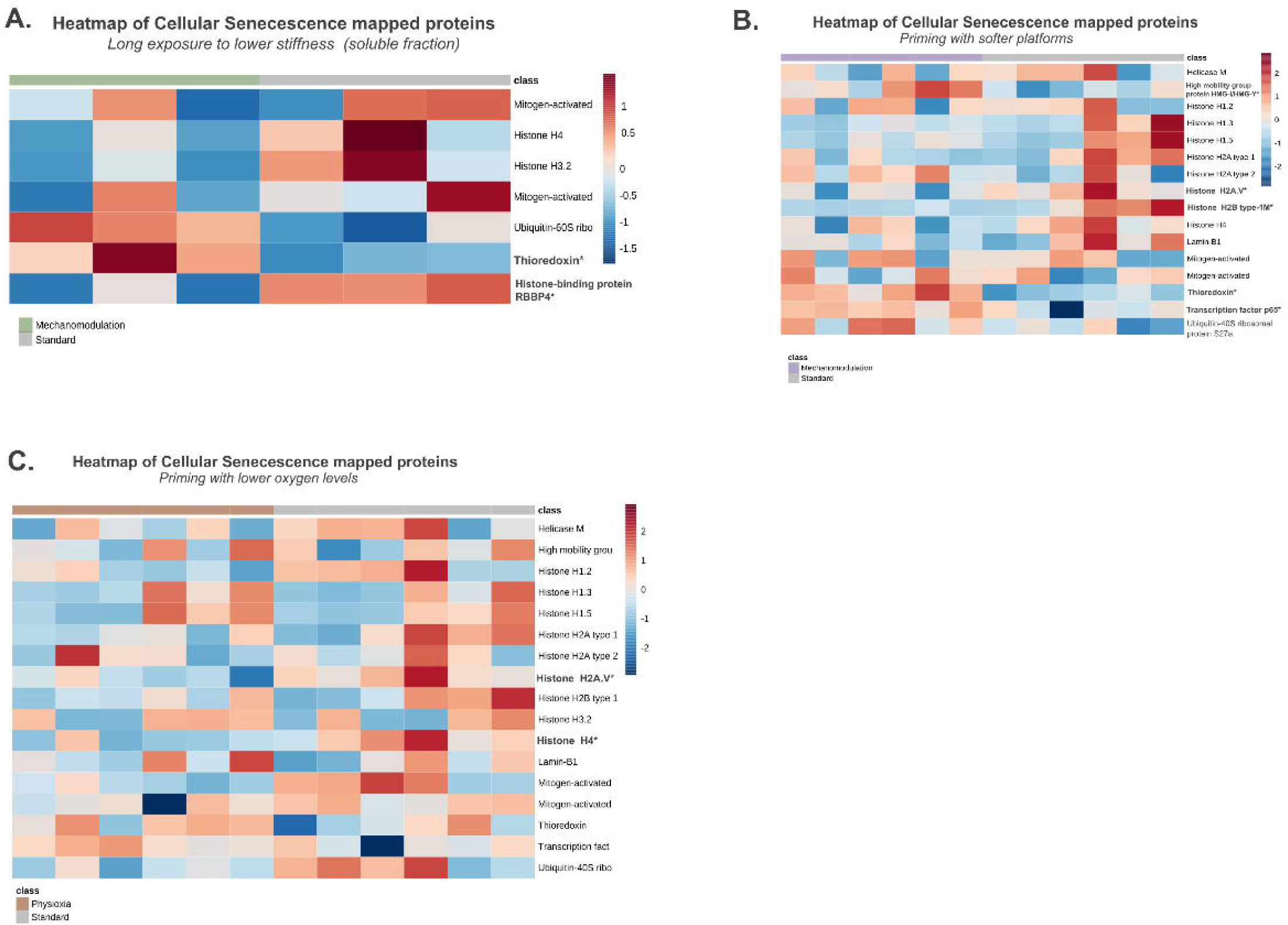
*Profiling of cellular senescence-associated proteins.* Heatmaps represent the levels of altered proteins associated with the cellular senescence of Reactome on long (membrane fraction) and priming experiments. Proteins highlighted in bold with (*) are statistically relevant (Kruskal-Wallis and t-test were applied on priming and long-exposure experiments, respectively; *p<0.05).

**Supplementary Table 1.**
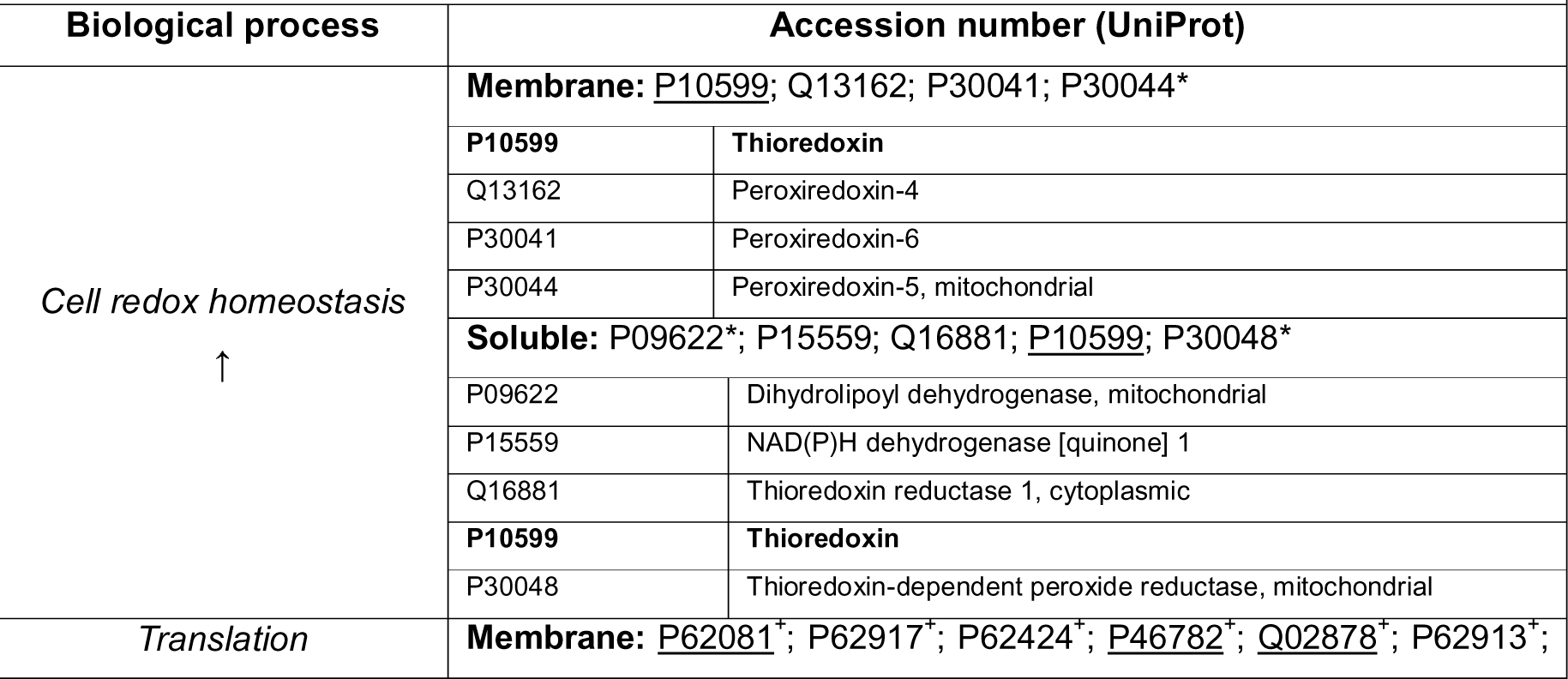
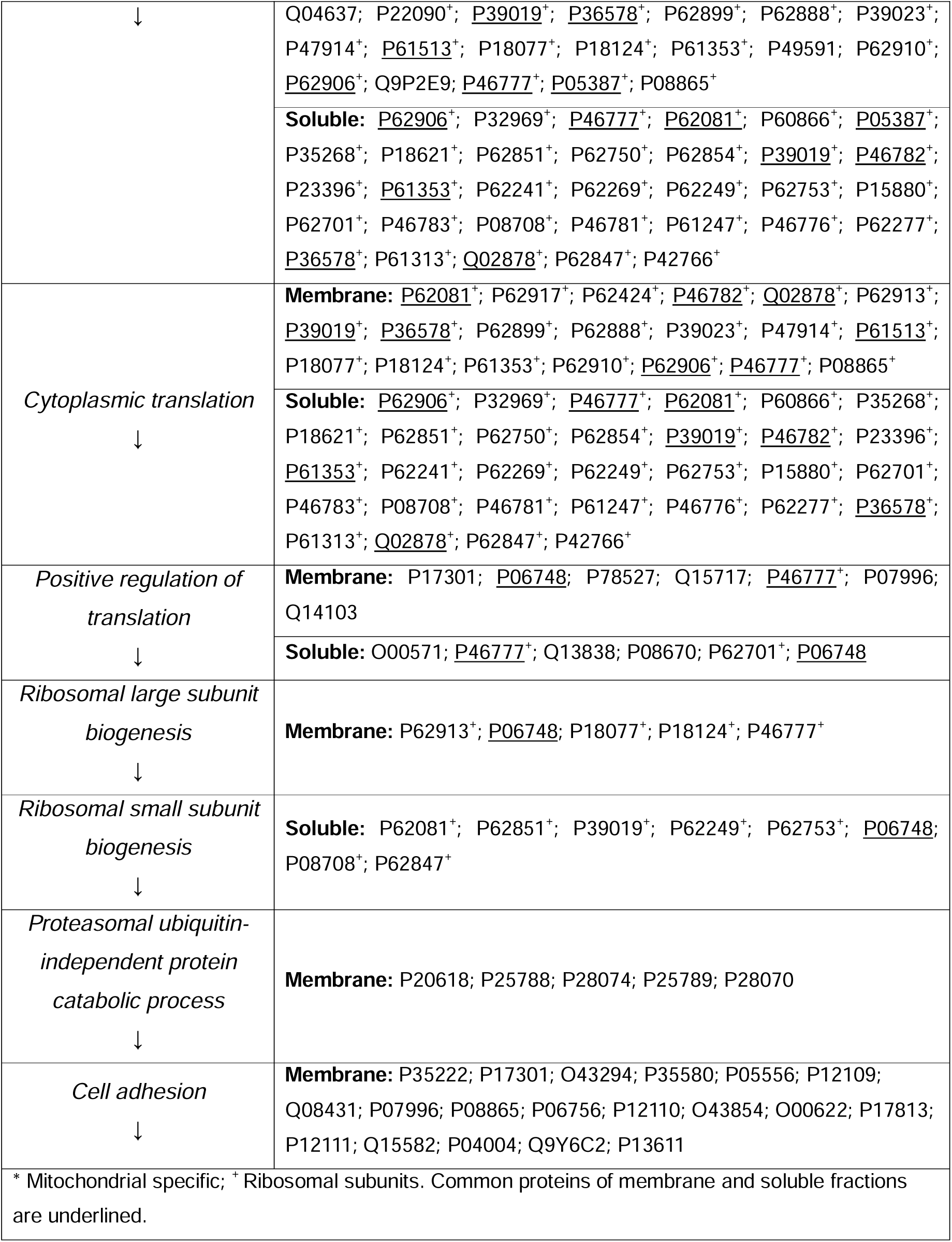
Proteins associated with significant biological processes when cultured on soft stiffness platforms.

**Supplementary Table 2.**
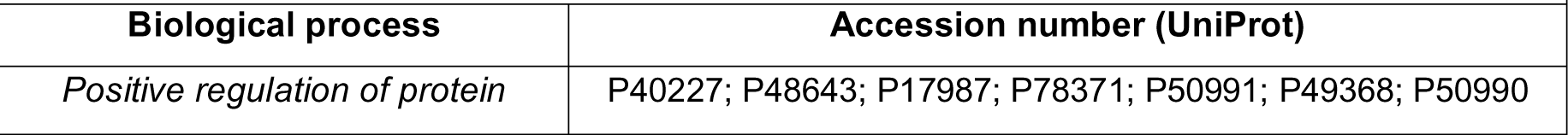
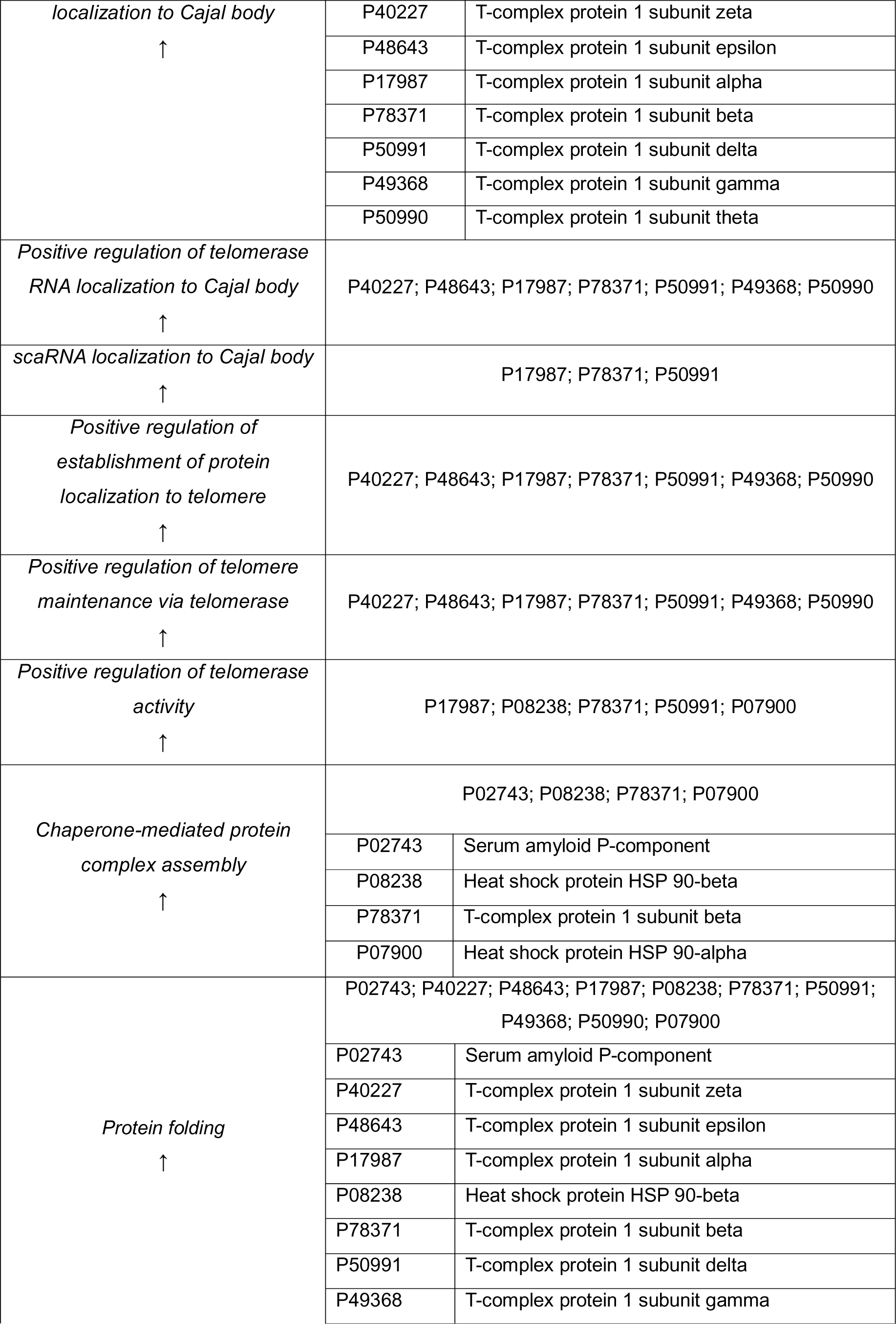
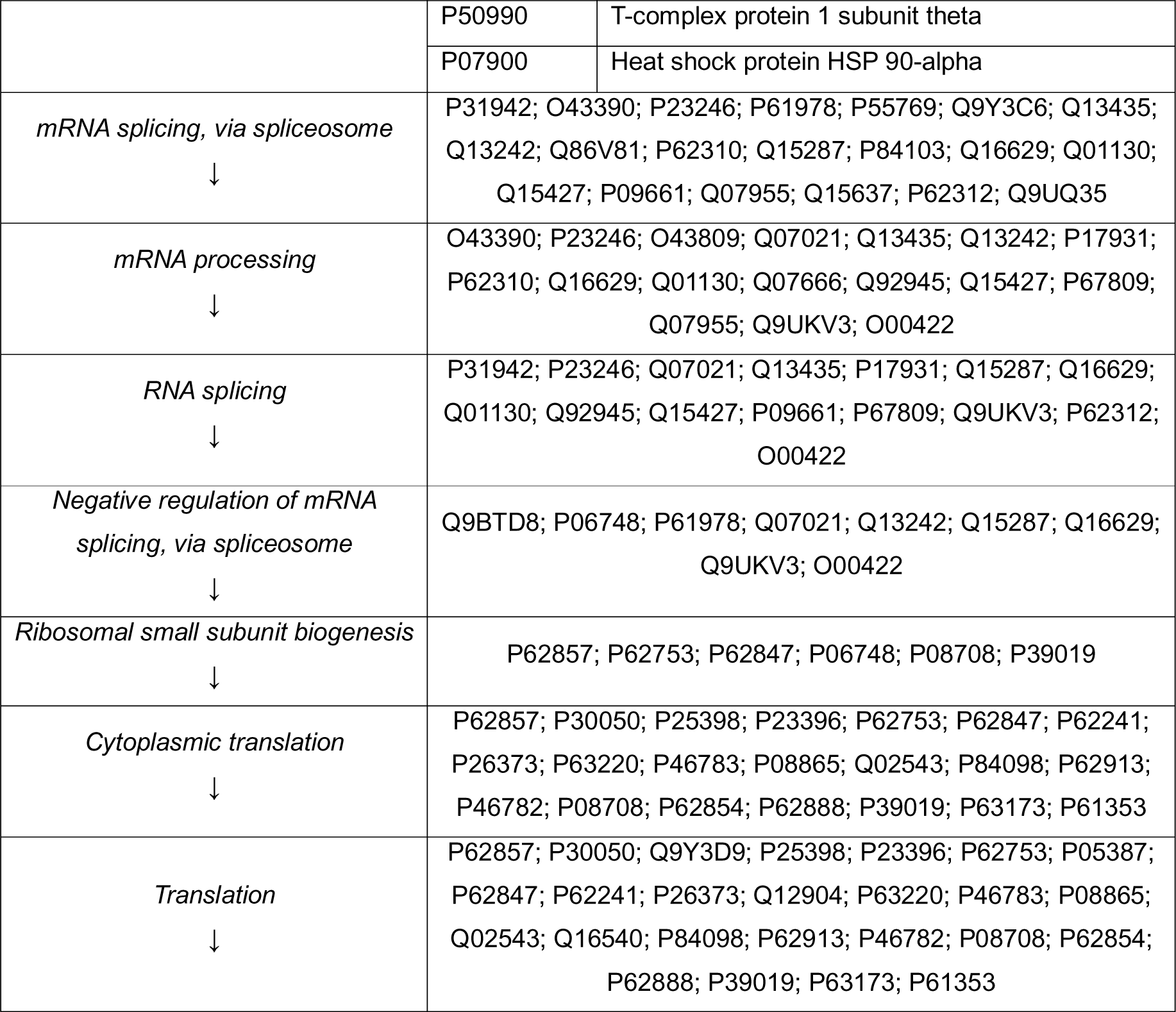
Proteins associated with significant biological processes when cultured under physiological levels of oxygen.

| Biological process | Accession number (UniProt) |
| --- | --- |
| *Positive regulation of protein* | P40227; P48643; P17987; P78371; P50991; P49368; P50990 |

|  |  |  |
| --- | --- | --- |
| *localization to Cajal body*<br>↑ | P40227 | T-complex protein 1 subunit zeta |
|  | P48643 | T-complex protein 1 subunit epsilon |
|  | P17987 | T-complex protein 1 subunit alpha |
|  | P78371 | T-complex protein 1 subunit beta |
|  | P50991 | T-complex protein 1 subunit delta |
|  | P49368 | T-complex protein 1 subunit gamma |
|  | P50990 | T-complex protein 1 subunit theta |
| *Positive regulation of telomerase RNA localization to Cajal body*<br>↑ | P40227; P48643; P17987; P78371; P50991; P49368; P50990 |  |
| *scaRNA localization to Cajal body*<br>↑ | P17987; P78371; P50991 |  |
| *Positive regulation of establishment of protein localization to telomere*<br>↑ | P40227; P48643; P17987; P78371; P50991; P49368; P50990 |  |
| *Positive regulation of telomere maintenance via telomerase*<br>↑ | P40227; P48643; P17987; P78371; P50991; P49368; P50990 |  |
| *Positive regulation of telomerase activity*<br>↑ | P17987; P08238; P78371; P50991; P07900 |  |
| *Chaperone-mediated protein complex assembly*<br>↑ | P02743; P08238; P78371; P07900 |  |
|  | P02743 | Serum amyloid P-component |
|  | P08238 | Heat shock protein HSP 90-beta |
|  | P78371 | T-complex protein 1 subunit beta |
|  | P07900 | Heat shock protein HSP 90-alpha |
| *Protein folding*<br>↑ | P02743; P40227; P48643; P17987; P08238; P78371; P50991; P49368; P50990; P07900 |  |
|  | P02743 | Serum amyloid P-component |
|  | P40227 | T-complex protein 1 subunit zeta |
|  | P48643 | T-complex protein 1 subunit epsilon |
|  | P17987 | T-complex protein 1 subunit alpha |
|  | P08238 | Heat shock protein HSP 90-beta |
|  | P78371 | T-complex protein 1 subunit beta |
|  | P50991 | T-complex protein 1 subunit delta |
|  | P49368 | T-complex protein 1 subunit gamma |
|  | P50990 | T-complex protein 1 subunit theta |
|  | P07900 | Heat shock protein HSP 90-alpha |
| *mRNA splicing, via spliceosome*<br>↓ | P31942; O43390; P23246; P61978; P55769; Q9Y3C6; Q13435; Q13242; Q86V81; P62310; Q15287; P84103; Q16629; Q01130; Q15427; P09661; Q07955; Q15637; P62312; Q9UQ35 |  |
| *mRNA processing*<br>↓ | O43390; P23246; O43809; Q07021; Q13435; Q13242; P17931; P62310; Q16629; Q01130; Q07666; Q92945; Q15427; P67809; Q07955; Q9UKV3; O00422 |  |
| *RNA splicing*<br>↓ | P31942; P23246; Q07021; Q13435; P17931; Q15287; Q16629; Q01130; Q92945; Q15427; P09661; P67809; Q9UKV3; P62312; O00422 |  |
| *Negative regulation of mRNA splicing, via spliceosome*<br>↓ | Q9BTD8; P06748; P61978; Q07021; Q13242; Q15287; Q16629; Q9UKV3; O00422 |  |
| *Ribosomal small subunit biogenesis*<br>↓ | P62857; P62753; P62847; P06748; P08708; P39019 |  |
| *Cytoplasmic translation*<br>↓ | P62857; P30050; P25398; P23396; P62753; P62847; P62241; P26373; P63220; P46783; P08865; Q02543; P84098; P62913; P46782; P08708; P62854; P62888; P39019; P63173; P61353 |  |
| *Translation*<br>↓ | P62857; P30050; Q9Y3D9; P25398; P23396; P62753; P05387; P62847; P62241; P26373; Q12904; P63220; P46783; P08865; Q02543; Q16540; P84098; P62913; P46782; P08708; P62854; P62888; P39019; P63173; P61353 |  |

**Supplementary Table 3.** Common proteins associated with translation on both mechanomodulated and physioxia culture conditions.

|  |  |  |
| --- | --- | --- |
| Biological process | Accession number (UniProt) | Name |
| Translation | P23396 | 40S ribosomal protein S3 |
|  | P62753 | 40S ribosomal protein S6 |
|  | P05387 | 60S acidic ribossomal protein P2 |
|  | P62847 | 40S ribosomal protein S24 |
|  | P62241 | 40S ribosomal protein S8 |
|  | P46783 | 40S ribosomal protein S10 |
|  | P08865 | 40S ribosomal protein SA |
|  | P62913 | 60S ribosomal protein L11 |
|  | P46782 | 40S ribosomal protein S5 |
|  | P08708 | 40S ribosomal protein S17 |
|  | P62854 | 40S ribosomal protein S26 |
|  | P62888 | 60S ribosomal protein L30 |
|  | P39019 | 40S ribosomal protein S19 |
|  | **P61353** | **60S ribosomal protein L27** |

**Supplementary Table 4.** Common proteins altered upon physiological priming.

|  |  |  |
| --- | --- | --- |
| Comparison | Accession number<br>(UniProt) | Name |
| ↑ *Mechanomodulated*<br>vs.<br>↑ *Physioxia* | P02794 | Ferritin heavy chain |
|  | P30740 | Leukocyte elastase inhibitor |
|  | Q9Y570 | Protein phosphatase methylesterase 1 |
| ↓ *Mechanomodulated*<br>vs.<br>↓ *Physioxia* | P24666 | Low molecular weight phosphotyrosine protein phosphatase |
|  | P46939 | Utrophin |
|  | Q9NRX2 | 39S ribosomal protein L17, mitochondrial |
|  | **P13611** | **Versican core protein** |
|  | O60462 | Neuropilin-2 |
| The highlighted proteins are involved in the relevant biological processes identified in long exposure and priming experiments, such as cell adhesion. |  |  |

**Supplementary Table 5.** Common proteins altered by mechanomodulation stimulus on long and short periods (priming)

|  |  |  |
| --- | --- | --- |
| Comparison | Accession number<br>(UniProt) | Name |
| ↑ *Long exposure*<br>Vs.<br>↑ *Priming* | P02794 | Ferritin heavy chain |
|  | Q16881 | Thioredoxin reductase 1, cytoplasmic |
|  | **P10599** | **Thioredoxin** |
| ↑ *Long exposure*<br>Vs.<br>↓ *Priming* | P08123 | Collagen alpha-2(I) chain |
| ↓ *Long exposure*<br>Vs.<br>↑ *Priming* | P07996 | Thrombospondin-1 |
|  | Q14157 | Ubiquitin-associated protein 2-like |
| ↓ *Long exposure*<br>Vs.<br>↓ *Priming* | P08708 | 40S ribosomal protein S17 |
|  | P02461 | Collagen alpha-1(III) chain |
|  | Q00341 | Vigilin |
|  | **P12110** | **Collagen alpha-2(VI) chain** |
|  | Q9Y6C2 | EMILIN-1 |
|  | P12109 | Collagen alpha-1(VI) chain |
|  | Q15582 | Transforming growth factor-beta-induced protein ig-h3 |
|  | Q07954 | Prolow-density lipoprotein receptor-related protein 1 |
|  | P12111 | Collagen alpha-3(VI) chain |
|  | O00622 | CCN family member 1 |
|  | **P13611** | **Versican core protein** |
| The highlighted proteins are involved in the relevant biological processes identified in long exposure and priming experiments, such as cell adhesion and cell redox homeostasis. |  |  |

**Supplementary Table 6.** Common proteins altered by controlled oxygen levels on long and short periods (priming)

|  |  |  |
| --- | --- | --- |
| Comparison | Accession number (UniProt) | Name |
| ↑ *Long exposure*<br>Vs.<br>↑ *Priming* | - | - |
| ↑ *Long exposure*<br>Vs.<br>↓ *Priming* | Q16658 | Fascin |
|  | P31930 | Cytochrome b-c1 complex subunit 1, mitochondrial |
| ↓ *Long exposure*<br>Vs.<br>↑ *Priming* | P05387 | 60S acidic 34ribosomal protein P2 |
|  | P02794 | Ferritin heavy chain |
| ↓ *Long exposure*<br>Vs.<br>↓ *Priming* | P24666 | Low molecular weight phosphotyrosine protein phosphatase |
|  | O75348 | V-type proton ATPase subunit G 1 |
|  | **P61353** | **60S ribosomal protein L27** |
|  | **P13611** | **Versican core protein** |
|  | P16403 | Histone H1.2 |
|  | P80723 | Brain acid soluble protein 1 |
|  | Q9NX24 | H/ACA ribonucleoprotein complex subunit 2 |
| The highlighted proteins are involved in the relevant biological processes identified in long exposure and priming experiments, such as cell adhesion and translation. |  |  |

**Supplementary Table 7.**
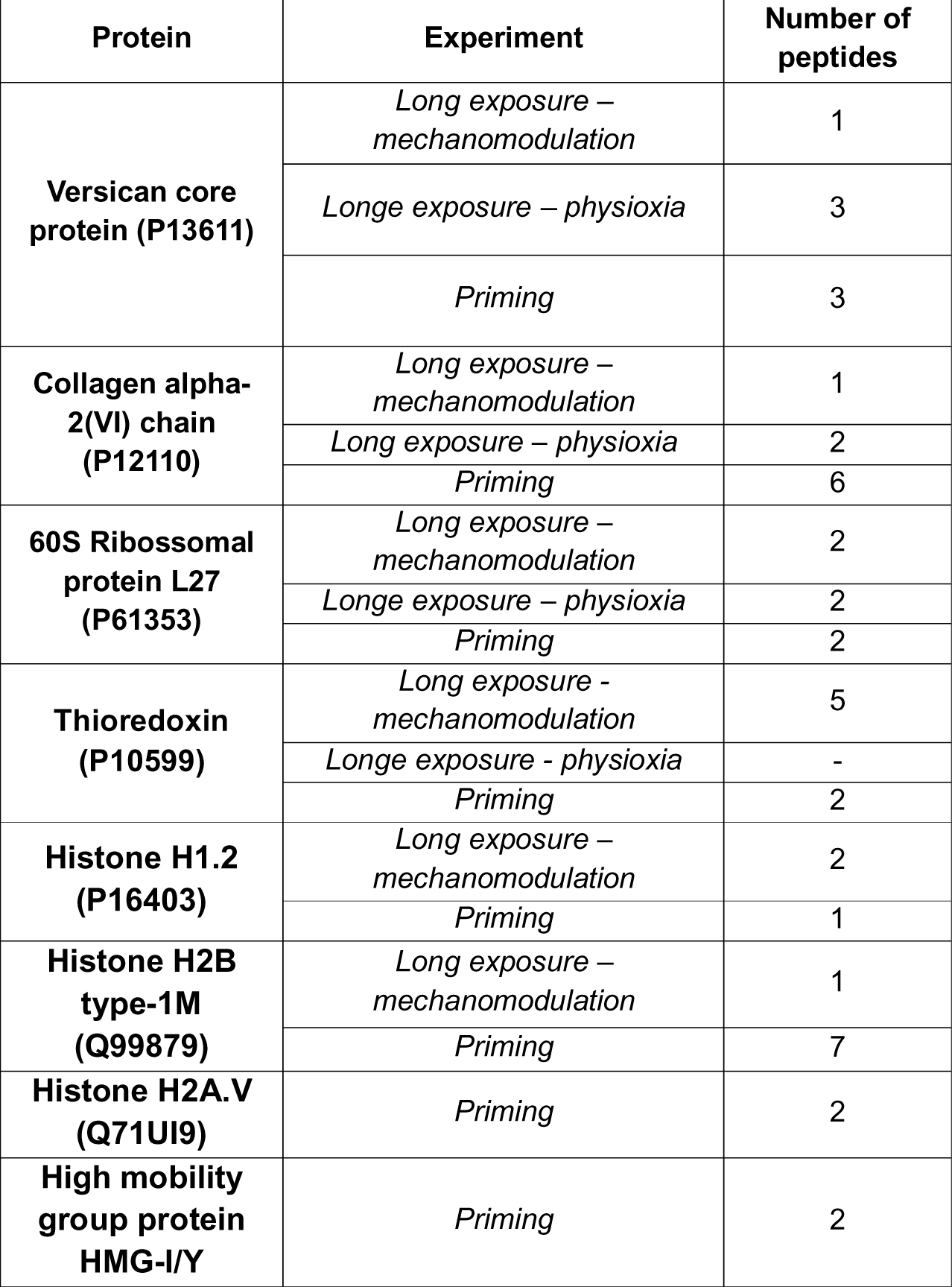
Number of peptides quantified for each highlighted protein (proteins in bold) in the different proteomic screenings.

**Supplementary Table 8.**
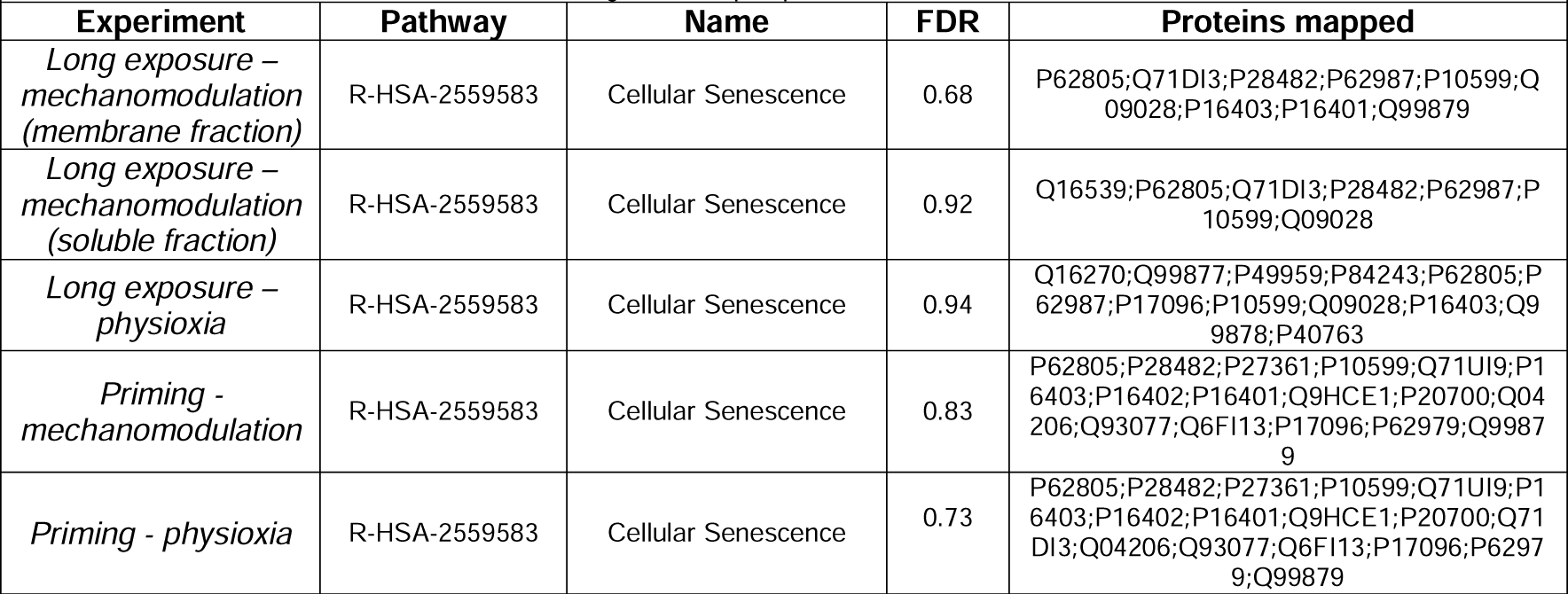
Mapped entities associated to the Reactome pathway “Cellular senescence” and “Transcriptional regulation of pluripotent stem cells”.

## Notes

### Competing Interest Statement

The authors have declared no competing interest.

