## Supplementary material for "Mimicking physiological stiffness or oxygen levels in vitro reorganizes mesenchymal stem cells machinery toward a more naïve phenotype": Methods in detail

### *Cell culture*

#### Isolation and expansion of UC-MSCs

All the procedures were approved by the Ethics Committee of the Faculty of Medicine, University of Coimbra, Portugal (ref. CE-075/2019) and were previously reported by our team (24). Succinctly, human UCs were obtained from healthy donors upon informed consent from the parent(s). Fragments from the UC matrix (Wharton’s jelly) were placed on a Corning® tissue-culture polystyrene (TCP) dishes with proliferation medium [Minimum essential Medium-α (MEM-α) (Gibco™) supplemented with 10% (v/v) MSC-qualified fetal bovine serum (FBS) (Hyclone, GE Healthcare) (only for long-term mechanomodulated UC-MSCs) or 5% (v/v) fibrinogen depleted human platelet lysate (HPL) (UltraGRO™, Helios) and antibiotics: 100 U/mL of Penicillin, 100 μg/ml Streptomycin and 2.5 μg/ml Amphotericin B or Antibiotic-Antimycotic (1×) (all from Gibco™)). Plates were incubated at 37ºC, with 5% CO_2,_ on a humidified incubator until colonies were observed by phase-contrast microscopy (approximately 10-14 days). Next, fragments were removed, the adherent cells were washed twice with sterile Phosphate Buffered Saline (PBS) and detached by adding 0.05% Trypsin-EDTA solution (Gibco™) for five minutes at 37°C on a humidified incubator. Then, detached cells were resuspended with washing medium (identical to proliferation medium but supplemented with heat-inactivated FBS (Gibco™)], and centrifuged for 5 minutes at 290×g at room temperature (RT). The supernatant was discarded, and the cell pellet was resuspended in the proliferation medium and re-seeded on a new plate (passage 1 – P1). Cells were left to proliferate until they reached 80-90% confluence. The procedure described above (after fragment removal) was repeated at each cell passage.

#### Expansion of UC-MSCs under controlled oxygen levels

For long exposure experiments, UC-MSCs were initially seeded at 4000/cm^2^ (P2) on a TCP culture dish, and placed on a humidified incubator (Binder), at 37ºC, with 5% CO_2_ – standard culture conditions - or in an InvivO₂® 400 Physoxia Workstation (Bakerruskinn), with 5% O_2_,5% CO_2_ and humidified environment - physioxia. To avoid any oxygen levels fluctuations, all procedures were performed within the workstation and all reagents used were left to equilibrate for at least 24 hours. When 80-90% confluent, cells were split 1:3 until P4.

For priming experiments, MSCs were kept at standard culture conditions until P3. Then, cells were seeded at 10,000/cm^2^ (P4) and left at standard conditions or physioxia for 48 hours.

#### Expansion of MSCs on soft platforms

Soft polydimethylsiloxane (PDMS) platforms were prepared as previously described by Domingues et al. (24). Briefly, the silicone elastomer base with the curing agent (Sylgard® 184, DOWSIL™) on a 40:1 proportion and cured for 4h at 80°C in an incubator (Memmert). Since this polymer is highly hydrophobic, its surface was chemically treated to facilitate protein coating and consequently cell adhesion, as described in previous studies (25). To do that, a solution of dd H_2_O/H_2_O_2_/HCl in a volumetric proportion of 5:1:1 for 5 minutes at RT generates silanol groups on the surface and turns the surface more hydrophilic. After washing three times with ddH_2_O, a solution of 10% (v/v) of 3-aminopropyltrimethoxisilane (3-APTMS, Alfa Aesar) in 96% ethanol (VWR) was added for 30 min, at RT. Next, substrates were washed abundantly three times (10 min each) with dd H_2_O with agitation, and 3% glutaraldehyde in PBS was added for 20 min, at RT, to promote covalent binding of coating proteins to the PDMS surface. Finally, PDMS substrates were washed three times with dd H_2_O with agitation and sterilized by exposure to ultraviolet light for 30 min in an air flow cabinet.

PDMS platforms were then incubated for 4 hours, at 37ºC, with a coating solution containing 10μg/mL of human plasma purified fibronectin (FN) (Merck Millipore) and 5μg/ml or 17μg/mL of rat tail type-I collagen (COL-I) (BD Biosciences or Millipore), to a final proportion of 1.4μg/cm^2^ and 0.7μg/cm^2^ or 2.4μg/cm^2,^ respectively. Before cell seeding, PDMS was washed twice with sterile PBS for protein excess removal. MSCs were initially seeded at 4,000/cm^2^ (P2) for long exposure experiments at standard culture conditions or in PDMS soft platforms – mechanomodulation. When 80-90% confluence was achieved, cells were split 1:3 until P4. For priming experiments, cells were expanded until P3 at standard conditions, seeded at 10,000/cm^2^ (P4) on PDMS platforms or TCP dishes, and left for 48 hours.

### *Proteomic analysis*

#### Protein extraction and sample preparation

Protein extracts were collected on ice by the addition of Laemmli buffer (120 mM Tris-HCl pH 6.8, 4% (w/v) Sodium dodecyl sulfate (SDS) and 20% (v/v) glycerol) except for long-exposure mechanomodulated MSCs, where the proteome was fractioned into soluble and membrane fraction (as previously described (24)). The total amount of protein was quantified using Pierce^TM^ 660nm Protein Assay (Thermo Fischer Scientific) according to the manufacturer’s instructions. Then, samples were denatured for 5 min at 95°C and sonicated at 60% amplitude, with pulses of 3’’ on and 2’’ off for one minute. Then, sample buffer 6× (350mM Tris-HCl pH 6.8, 30% (v/v) glycerol, 10% (w/v) SDS, 0.93% (w/v) dithiothreitol (DTT)) was added to the sample to obtain a final concentration of 1×, and proteins were alkylated by the addition of 40% acrylamide to a final concentration of 1:15 (v/v). The same amount of a protein standard MBP-GFP was added to each protein extract (25). For protein identification, pools of each condition (70 μg) were prepared, while for protein quantification 50 μg of each sample was loaded individually on the gel. Proteins were resolved through a short-GeLC approach and stained with Coomassie Brilliant Blue G-250 (26, 27). Gel bands were then cut into five or three band, concerning protein identification or quantification, respectively, and destained using a 50 mM ammonium bicarbonate with 30% acetonitrile solution. Proteins were then digested overnight, in gel, by trypsin (0.01 mg/ml), and digested peptides were extracted by the addition of solutions with an increasing percentage of acetonitrile (30, 50 and 98%) and 1% formic acid.

### *Data analysis*

Each fraction of pooled samples was acquired individually in DIA mode. ProteinPilot™ software was used to generate the ion library of the precursor masses and fragment ions (v5.0, Sciex, Framingham, MA, USA), combining all files from the pools. The data was searched against the reviewed Human (Swissprot) database (downloaded on 21^st^ October 2021), setting cysteine alkylation by acrylamide, digestion by trypsin, and gel-based ID factors. The quality of the identifications was assessed by an independent False Discovery Rate (FDR) analysis, which was performed using the target-decoy approach provided by the software.

For protein relative quantification, samples were acquired on DIA mode/SWATH. Data was processed using the SWATH™ plug-in for PeakView™ (v2.0.01, Sciex®, Framingham), setting 15 peptides per protein, and 5 transitions per peptide. Only peptides present in at least 1/3 of the samples per condition with an FDR below 1% were quantified. Data was normalized to the total intensity.

### *Measurent of oxygen levels*

The levels of oxygen dissolved in the culture media were continuously monitored on a MW96, using Resipher (Lucid Scientific), either in a standard humidified incubator or in the InvivO₂® 400 Physoxia Workstation. Data was acquired in duplicates or triplicates for each experimental condition, and oxygen levels were measured on each well with or without UC-MSCs seeded.

### *Proliferation kinetics*

UC-MSCs were expanded as previously described. Before seeding, the number of cells was manually calculated using a hemocytometer. The population doubling (PD) was determined at each passage using the following formula: $PD=[({log}_{10}\left( N_{H} \right)-{log}_{10}\left( N_{I} \right)]/{log}_{10}\left( 2 \right)$, where $N_{H}$ is the number of harvested cells and $N_{I}$ is the initial number of plated cells. Also, the cumulative total number of cells (TNC) at each passage was assessed using the formula $TNC=N_{H}\times B/N_{I}$, where B is the total number of cells of the previous passage.

### *Senescence β-Galactosidase Staining*

Cell senescence was assessed by the β-Galactosidase staining kit (#9860, Cell Signalling). The protocol was in agreement with the manufacturer's instructions. Birefly, cells were washed with PBS fixed with the fixative solution provided, and the β-Galactosidase staining solution was added. Then, UC-MSCs were incubated overnight at 37ºC on a dry incubator without CO_2_. On the following day, images were captured on a contrast phase microscope (Axiovert 40C), using the ZEN Microscopy Software (3.3), to identify the development of a blue color.

*Western blot*

Samples were prepared as previously described for the proteomics screening. Protein extracts (12.5µg) were loaded and resolved by using SDS-PAGE with either 10% or 12.5% (w/v) acrylamide–bisacrylamide gels (Bio-Rad). Subsequently, proteins were transferred to a low fluorescence polyvinylidene fluoride (PVDF) membranes using a Trans-Blot Turbo Transfer System (BioRad) for 30 min at a constant voltage of 25 V (up to 1 A). Membranes and solutions used were provided with the transfer kit (TBT RTA TRANSFER KIT, Bio-Rad). Membranes were stained with Ponceau S staining Solution (Thermo Scientific) to confirm protein transfer. After removing the stain, membranes were blocked for 1 hour with 5% (w/v) skimmed milk powder in TBS-Tween 20 (Biorad) (TBS-T) [0.1% (v/v)]. Then, membranes were incubated with primary antibodies overnight at 4ºC followed by 1 hour at room temperature, with agitation (YAP 1:1000 (#101199, Santa Cruz); CTGF 1:500 (#365970, Santa Cruz); cofilin-1 1:1000 (#51755, Cell Signalling), phospho-cofilin 1µg/ml (#44-1072G, Invitrogen)). Following this incubation, membranes were washed three times (5 minutes, with agitation) with TBS-T and incubated with the respective secondary antibody (anti-mouse 1:10,000 (115‐055‐003), anti-rabbit 1:20,000 (111‐055‐003)) with the respective secondary antibodies conjugated with alkaline phosphatase for 1 hour, at room temperature. Finally, membranes were incubated with enhanced chemifluorescence substrates (ECF – GE Healthcare) for 5 minutes and imaged using a VWR Imager (VWR). To determine the total amount of protein in each lane, membranes were stained with SERVA purple (SERVA electrophoresis, Enzo) following the manufacturer’s guidelines. Images were analyzed with Image Lab software, version 5.1 (BioRad).
